# Structural basis for lipid transfer by the ATG2A-ATG9A complex

**DOI:** 10.1101/2023.07.08.548186

**Authors:** Yang Wang, Selma Dahmane, Rujuan Ti, Xinyi Mai, Lizhe Zhu, Lars-Anders Carlson, Goran Stjepanovic

## Abstract

Autophagy is characterized by the formation of double-membrane vesicles called autophagosomes. ATG2A and ATG9A play an essential role in autophagy by mediating lipid transfer and re-equilibration between membranes for autophagosome formation. Here we report the cryo-EM structures of human ATG2A-WIPI4 complex at 3.2 Å, and ATG2A-WIPI4-ATG9A complex at 7 Å resolution. The ATG2A structure is characterized by a central hydrophobic cavity formed by a network of β-strands that facilitates lipid transfer, and highly flexible N- and C-terminal domains. Molecular dynamics simulations of the ATG2A N-terminal domain revealed the mechanism of lipid-extraction from the donor membranes while the ATG2A-ATG9A complex structure provides insights into the later stages of the lipid transfer reaction. ATG9A-ATG2A structural analysis revealed a 1:1 stoichiometry, directly aligning the ATG9A lateral pore with ATG2A lipid transfer cavity, hence allowing for a direct transfer of lipids from ATG2A. The ATG9A trimer can interact with both N- and C-terminal tip of rod-shaped ATG2A. Cryo-electron tomography of ATG2A-liposome binding states shows that ATG2A tethers lipid vesicles at different orientations. In summary, this study provides a molecular basis for the growth of the phagophore membrane, and lends structural insights into spatially coupled lipid transport and re-equilibration during autophagosome formation.

## Main

Autophagy is a highly conserved cellular process from yeast to humans, which is important to maintain cellular homeostasis [1]. Autophagy recycles nutrients to resist stress and survive by degradation of the cell’s own damaged organelles, aggregated proteins and other macromolecules [2, 3]. Autophagy plays a key role in both physiology and pathology, its dysregulation leads to different diseases, including cancer, neurodegeneration, cardiovascular disease, diabetes and infections [4, 5]. Now, more than 40 proteins have been identified participating in autophagy regulation [6, 7]. The formation of autophagosome involves three steps, (i) nucleation, it starts at a double membrane structure, named phagophore; (ii) elongation, the phagophore membrane growth; (iii) phagophore closure and maturation [8]. The autophagosome ultimately fuses with lysosome forming autolysosome to degrade the cargo and recycle [3]. Autophagosome maturation takes ∼10 minutes and requires a lipid transfer rate of ∼5,000 lipids per second [9–12]. Recent research reported that period from phagophore initiation to autophagosome fusion with lysosome only takes ∼175 s in U2OS cells [13]. Autophagy related (ATG) proteins act sequentially to control the autophagosome biogenesis. During the nucleation and elongation process, membrane tethering and lipids transport is necessary for phagophore membrane growth, and the core machinery responsible for this process is ATG2A-WIPI4 -ATG9 complex. ATG2A is a peripheral membrane protein that tethers membranes and transports lipids from donor membranes such as ER to phagophore with the help of WIPI proteins [14–17]. ATG2A has a rod-shaped structure with a N-terminal Chorein_N-like domain (Chorein_N), central hydrophobic cavity suggested to accommodate and transfer lipids, and the C-terminal localization region (CLR) followed by autophagy-related protein C-terminal domain (ATG_C). In yeast Atg2, the N-terminal domain is required for association with the ER [15] while the C-terminal region binds to pre-autophagosomal membranes [15, 18]. Similarly, C-termina CLR domain is required for the localization of ATG2A to the phagophore by amphipathic α-helices [19]. As a result, ATG2 may tether these membranes together at the ER-phagophore contact sites to catalyze lipid transfer and drive phagophore membrane expansion [20, 21]. ATG_C is dispensable for autophagy and involved in ATG2 localization to lipid droplets [19]. WIPI4 (Atg18/WD-repeat protein interacting with phosphoinositides, Atg18 in yeast and WIPI1-4 in mammals) is recruited by phosphatidylinositol 3-phosphate (PI3P) enriched in phagophore. WIPI4 contributes to ATG2A anchoring to PI3P-positive phagophore membrane and lipid transfer between phagophores and ER [17, 21, 22]. In addition to WIPI4, ATG2A can interact with GABARAP (gamma-aminobutyric acid receptor-associated proteins; mammalian homologs of yeast Atg8) through LIR sequence which is approximately 30 residues before WIPI4 interaction motif. GABARAP is bound to autophagic membranes through covalent conjugation with phosphatidylethanolamines (PE), and its interaction with ATG2A is essential for phagophore formation [14]. ATG9A is the sole trans-membrane protein in the autophagy pathway where it catalyzes lipid redistribution between leaflets, leading to the phagophore expansion [18, 23–25]. Golgi-derived small vesicles loaded with ATG9 (termed ATG9 vesicles) are shown to be the seed membrane for autophagosomes by initiating lipid transfer from donor compartments such as the ER, in yeast and mammals. [18, 24, 26–28]. Around 60–80% of the autophagosome membrane lipids are derived from lipid transfer or *de novo* synthesis during expansion [28]. ATG9-containing vesicles such as Golgi derived vesicles or ATG9A and ATG16L1 containing recycling endosomes can also serve as membrane sources for phagophore expansion [17, 29–31]. ATG9A is a homo-trimer with a central membrane-penetrating pore, which can open and close in two different states. The central pore is connected to the cytosol through the lateral hydrophilic cavity within each protomer [23, 24]. ATG9A funnel-like trimeric architecture allows for a nondirectional lipid flipping through connecting the central pore to the three lateral cavities. ATG9A can interact with ATG2A lipid transporter [32] and potentially directly accept and scramble incoming lipids, thereby correcting lipid asymmetry in the growing phagophore membrane. This model is supported by yeast studies showing that the Atg9–Atg2 association promotes the Atg18 recruitment, and collectively are important to establish ER-phagophore contact sites [18]. The molecular details of the ATG2A-ATG9A complex, their interaction interface, mechanism and functional consequences remain to be determined.

To gain further insights into ATG2A-WIPI4-ATG9A lipid transport mechanism, we combined cryo-EM and cryo-electron tomography (cryo-ET) to generate a structural model of human ATG2A and ATG9A-ATG2A-WIPI4 complex. Molecular dynamics simulations of ATG2A N-terminal domain revealed how lipids are extracted from donor membranes at the starting stage and transported through this domain, which was validated by *in vitro* activity assays. Structural analysis of the ATG9A-ATG2A-WIPI4 complex revealed that the ATG9A trimer can interact with both N- and C-terminal end of ATG2A through a close alignment of the ATG9A lateral pore and the ATG2A lipid transfer cavity orifice, allowing for a direct lipid handover. Taken together, we provide structural and mechanistic insights, and propose a model of ATG9A-ATG2A-WIPI4 mediated lipid transport during autophagosome formation.

## Results

### The Cryo-EM structure of human ATG2A-WIPI4 complex

Human ATG2A is a 213 kDa protein which functions as lipid transporter in autophagosome biogenies. To understand the structural details of ATG2A, we reconstituted ATG2A, WIPI4 and ATG2A-WIPI4 complex purified by affinity resin and size exclusion chromatography **(Fig. S1)**. Cryo-EM images were collected and processed as detailed in the Methods section **(Fig. S2)**. The overall resolution for the ATG2A-WIPI4 structure is around 3.23 Å **(Fig. 1c, S2c)**. Most of the main chain in ATG2A could be traced except two long loop regions (304-424 and 1237-1416) that were not visible in the map **(Fig. 1a).** The N-terminal domain responsible for lipids extraction as well as ATG_C were also too flexible to directly model. ATG2A is about 180-200 Å in length and 30-50 Å in width with a hydrophobic central cavity **(Fig. 1b)**. The central cavity is formed by the association of 34 pairs of β-strands into twisted β-sheet that gradually become wider towards the C-terminus **(Fig. 1d-e)**. The cavity is not completely closed; the side opening is located opposite to the β-sheet lining of the cavity walls which spirals through the length of the molecule with a left-handed rotation of about 270° **(Fig. 1f-g)**. There are repeating patterns of the β-strands arrangement, 5 pairs of antiparallel β-strands form a repeating group with total of 6 repeats **(Fig. 1e)**, and commonly 1-2 residues link each pair of β-strands that allows formation of the cavity. Longer loops and α-helices intercede each β-strands group and some of the β-strands pairs, and partially occupy the side opening of the central cavity. The inner side of the cavity is mainly composed of hydrophobic sidechains which may facilitate the lipids acyl chain passthrough the channel **(Fig. 2c)**. The cavity length is 180-200 Å, and the width ranges from 24 Å at its C-terminus to 16 Å in the middle part of the protein, which is measured by the Cα-Cα in midpoint of each ꞵ-stand pair **(Fig. 2a)**. This would allow ATG2A to accommodate about 15 phospholipid molecules. ATG2A model is divided into three regions, N-terminal region (D173-S570), middle passage region (A577-L1128) and C-terminal region (A1129-D1735). The C-terminal portion of the central cavity is relatively wide and narrows down towards the middle passage region **(Fig. 2a)**. Middle passage region is characterized by several interacting hydrophobic residues: Leu671-Leu696 and Leu603-Leu694 forms hydrophobic interactions, and Phe678 points towards the center of the cavity and interacts with Leu603 **(Fig. 2b, Fig S2e)**. These interactions restrict central cavity diameter to ∼4Å, and may necessitate conformational changes in order to allow for lipid transfer through the protein. In order to gain insights in to the structure and location of ATG_C, which was unresolved in the cryo-EM structure, we performed chemical crosslinking combined with mass spectrometry (XL-MS). We used the MS-cleavable crosslinker DSBU (disuccinimidyl dibutyric urea), that connects primary amine groups of lysine residues within Cα-Cα distances up to ∼32 Å [33]. XL-MS analysis revealed several satisfied crosslinks within the ATG_C domain that were consistent with the AlphaFold helical bundle model, as well as crosslinks with Cα-Cα distances >32 Å that indicate alternate conformations in solution **(Fig. S3; Extended Table 1)**. ATG_C formed crosslinks with central parts of the ATG2A twisted β-sheet, including several lysine residues located at the edge of the lipid transfer cavity, at considerable distance from the C-terminus. Thus, it likely that ATG_C folds back to the ATG2 middle passage region where it may engage in transient interactions with the lipid transfer cavity. Collectively, XL-MS indicates that ATG_C is highly flexible relative to the rest of protein, explaining its absence from the final cryo-EM reconstruction and identifying potential long-range contacts with the β-sheet lining of the cavity walls **(Fig. S3b)**.

**Fig. 1:**
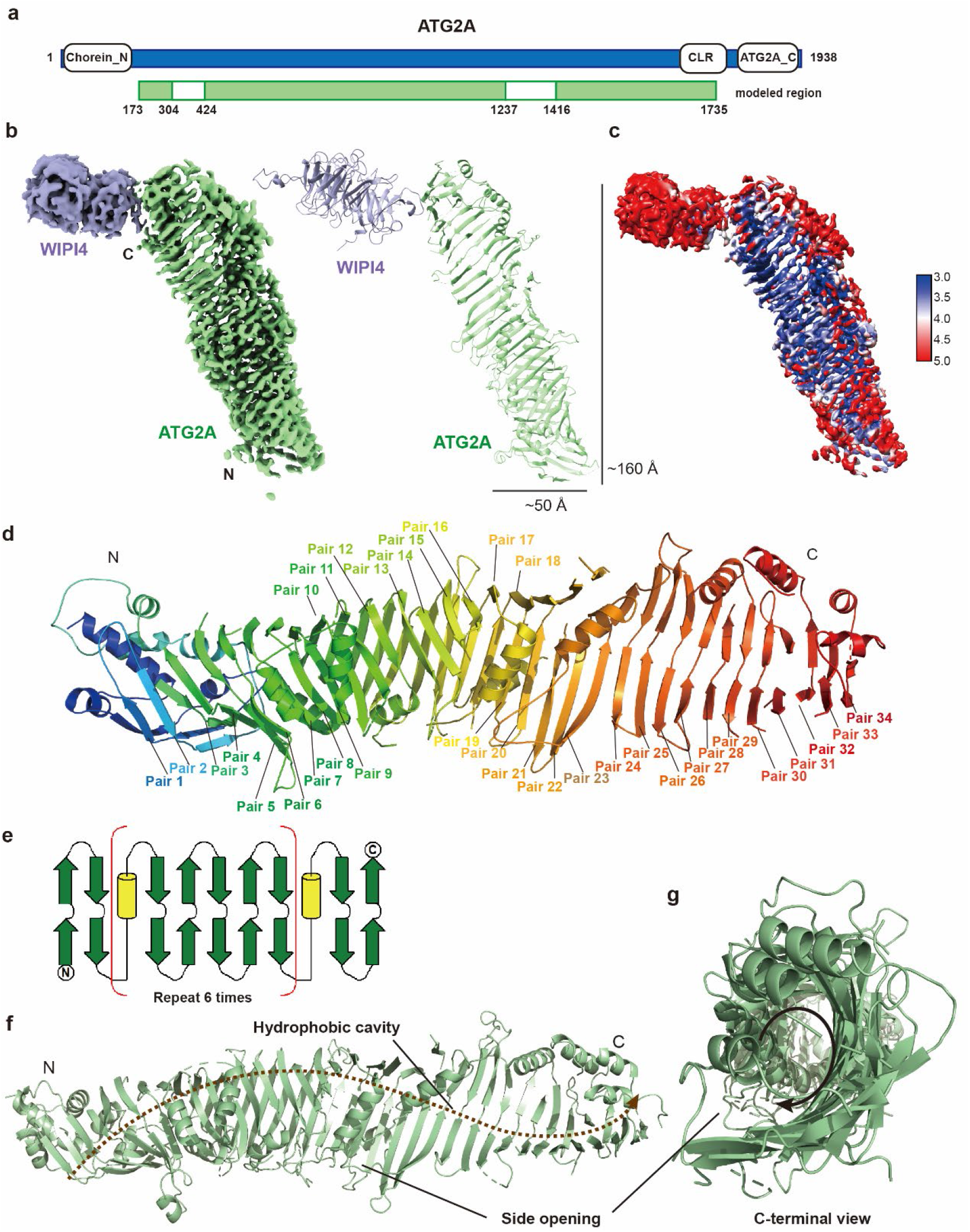
The cryo-EM structure of ATG2A. **a,** Domain organization diagram of human ATG2A with modelled regions are indicated in green. **b,** Depiction of density map and corresponding model of ATG2A-WIPI4 complex. ATG2A is colored in green and WIPI4 in purple. **c,** Depiction of local resolution of the ATG2A-WIPI4. **d,** ATG2A is composed of 34 pairs of β-stands forming the cavity. The first two pairs of β-stands are the flexible N-terminal region, which was generated based on the crystal structure of *sp*Atg2NR. **e,** The secondary structure diagram shows the arrangement of the 34 β-stands pairs. 5 pairs of antiparallel β-strands form a repeating group with total of 6 repeats. **f,** The cartoon shows ATG2A cavity is twisted with a side-opening from side view and C-terminal view **(g)**. The lipid may transfer through the hydrophobic channel in a spiral way as the arrow indicates.

**Fig. 2:**
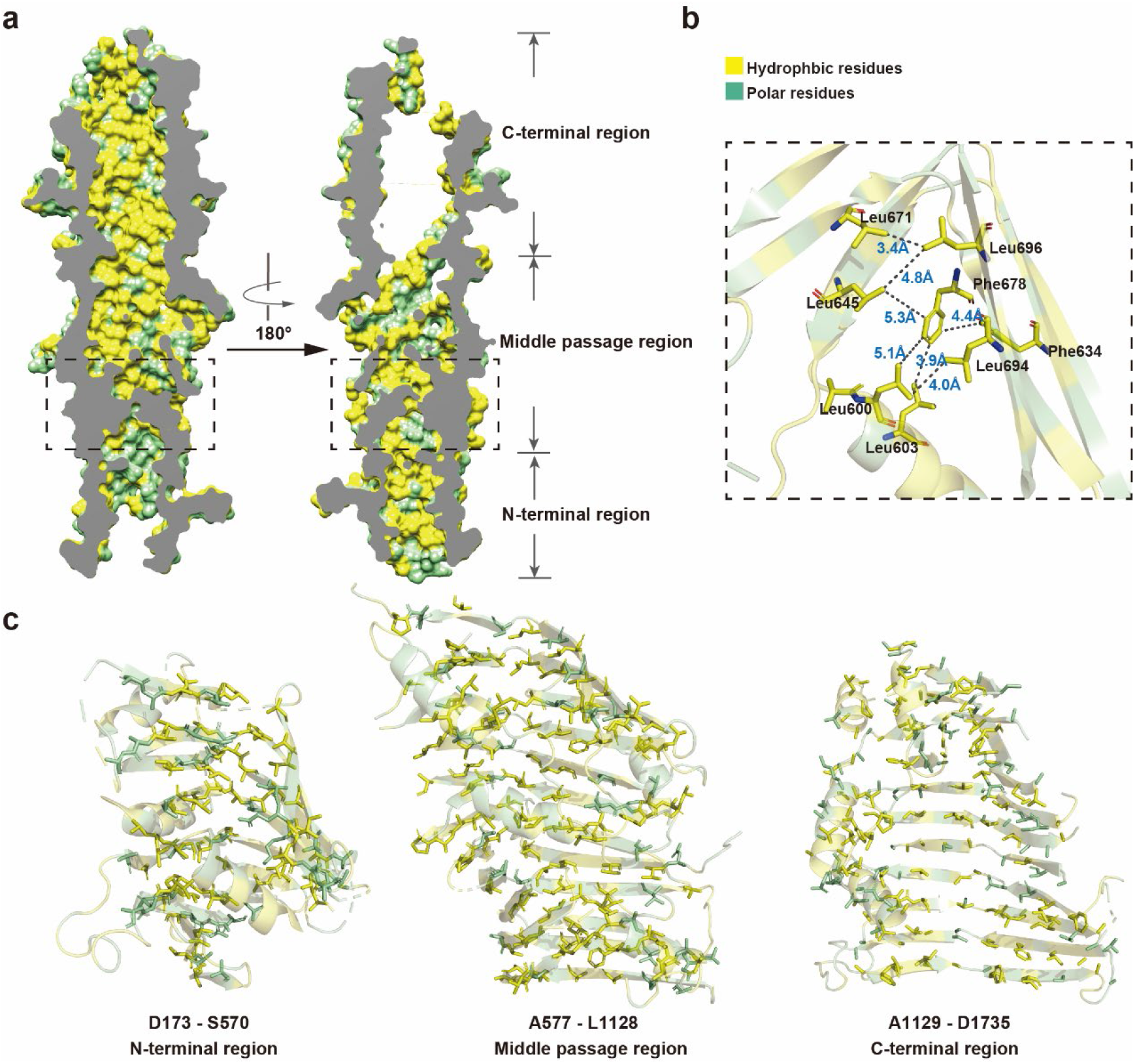
The structural details of ATG2A hydrophobic cavity. **a,** The cross sections of ATG2A model surface. We divided the model into three parts, N-terminal region, middle passage region and C-terminal region. The hydrophobic residues are colored in yellow, and polar residues are colored in green. **b,** The sidechain details of region A577 to D707 labeled in dash line box in **(a).** Distance between the hydrophobic residues is indicated. **c,** N-terminal region, middle region and C-terminal region are shown with the sidechains pointing toward cavity.

### Mechanism of lipid-transfer by N-terminal chorein domain

ATG2A N-terminal region (ATG2A^NR^) includes the chorein_N motif found in yeast Atg2, and several other lipid transfer proteins [34]. Yeast Atg2 N-terminal domain (Atg2^NR^) is responsible for ER anchoring [15], and capable of lipid extraction from membranes *in vitro* [35]. ATG2A^NR^ was too flexible to model and not clearly traceable in the density maps. To gain insights into such lipid extraction, we performed molecular dynamics (MD) simulations of *sp*Atg2^NR^ (PDB 6A9E) **(Fig. 3a)** alone and with DOPC micelles.

**Fig. 3:**
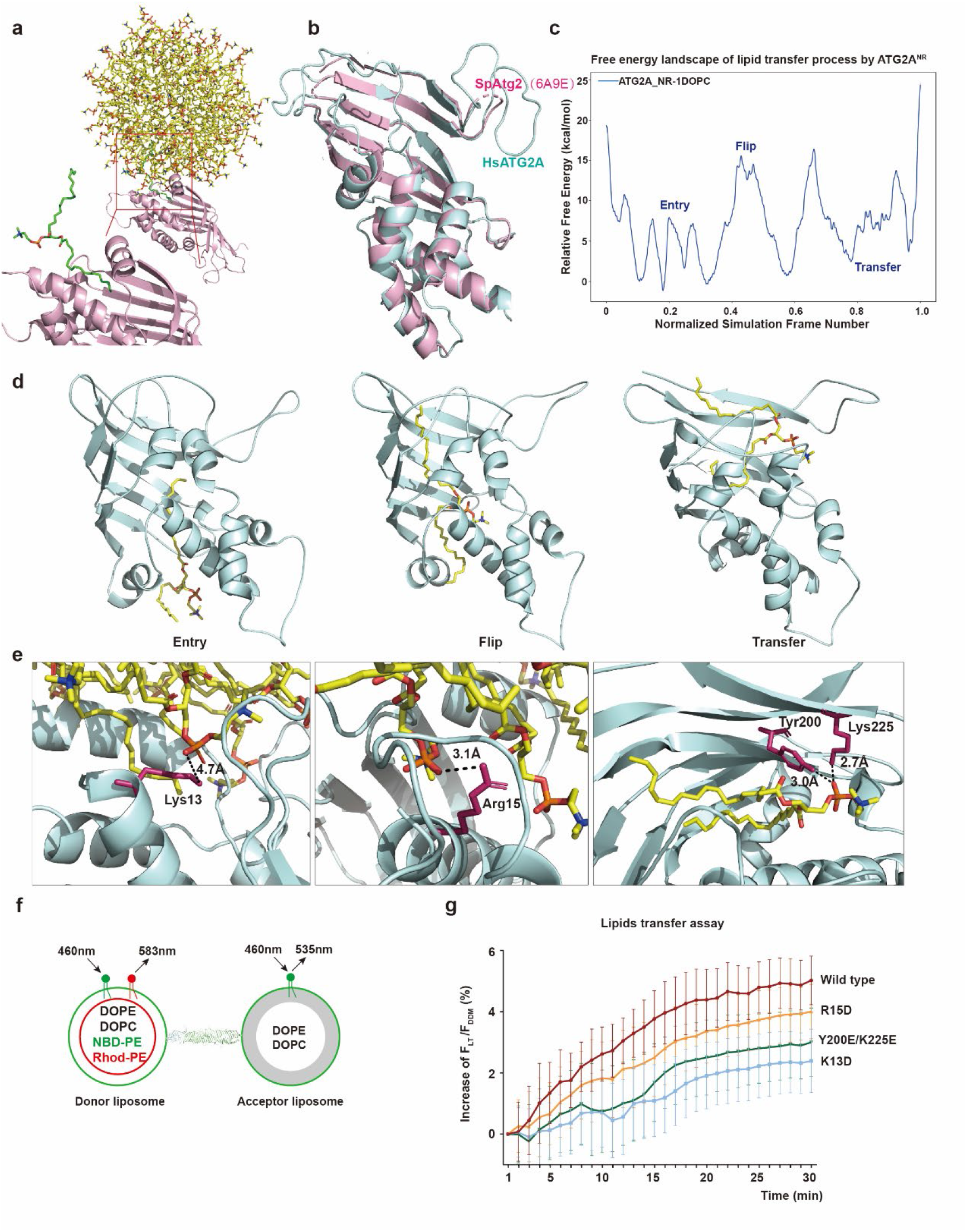
The simulation of lipids translocation process by ATG2A_NR_. **a,** The simulation of *sp*Atg2^NR^ (6A9E) [35] extracted lipids from DOPC micelle. *sp*Atg2 is colored in pink and the extracted lipid in green. **b,** The structural comparison of ATG2A^NR^ (8-236) and it’s simulation template *sp*Atg2 (6A9E). ATG2A^NR^ is colored in pale cyan and *sp*Atg2 in pink. **c,** The free energy landscape of ATG2A^NR^ during lipid transfer process. **d,** Three representative processes at the beginning of lipid transfer process including the entering (left), flipping (middle) and transporting (right). **e,** The key interaction sites between ATG2A^NR^ and lipids, Lys13 (top), Arg15 (middle), Tyr 200 and Lys225 (bottom). **f,** The activity assay model of ATG2A transfer lipids *in vitro*. **g,** The comparison of lipids transfer activity between wild type and mutants. The lipid transfer activity is measured by the NBD fluorescence which is not quenched by Rhodamine. F_LT_ means the NBD fluorescence intensity at each time point, followed by treatment with 0.4% DDM for 40 min and denoted as F_DDM_, these two values are measured at 535 nm. The increase in F_LT_/F_DDM_ was calculated by subtracting F_LT_/F_DDM_ of control group from protein-containing group, and minus the value at 1 min point. And the curve shows the mean values from three independent experiments.

Since the binding process between the protein and micelle is a random collision, we observed three different binding modes in three independent MD simulations. In two of the modes, the micelle forms contact with the lateral helix of *sp*Atg2^NR^ (**Fig. S4a**) or the edge of the folded sheet (**Fig. S4b**), where the lipid acyl chain in the micelle is far from the pocket. We therefore chose the third binding mode which facilitates lipid entry into the pocket to perform subsequent simulations **(Fig. 3a)**. Our sub-microsecond simulation had been sufficient to show that, when encountering a DOPC micelle, *sp*Atg2^NR^ is able to spontaneously absorb one phospholipid acyl chain to the entry of the lipid-transfer channel located on its N-terminal helical region **(Fig. 3a, Video 1)**.

We generated a homology model of ATG2A^NR^ (8-236 AA) based on the crystal structure of *sp*Atg2^NR^ **(Fig. 3b)** and modeled one DOPC in a similar position at the entry. MD simulations showed that ATG2A^NR^ can form stable complex with DOPC, similar to *sp*ATG2^NR^ **(Video 2)**. However, subsequent lipid-transfer through ATG2A^NR^ could occur at significantly longer timescale and is therefore beyond the reach of conventional MD simulations. To further dissect the starting stage of lipid-transfer by ATG2A^NR^, we located the minimum free energy path (MFEP) for this process via the recently matured travelling-salesman automated path searching (TAPS) algorithm [36]. Requiring minimal input information, TAPS is highly efficient and has successfully revealed high dimensional MFEPs for proteins with hundreds of residues [37–39]. Starting from the encounter DOPC-ATG2A^NR^ complex (without micelle or ATG9A/WIP4), we used biased sampling dragging the single lipid to penetrate the ATG2A^NR^ transfer-channel and generated an initial biased path for lipid-transfer. The MFEP was further obtained via a 100-ns TAPS optimization of this path **(Fig. S5a-b)**. We then used umbrella sampling [40] to calculate the free energy landscape along the MFEP (overlap of simulation windows shown in **Fig. S5c**). The high barrier of ∼15 kcal/mol on the MFEP **(Fig. 3c)** indicates that lipid-transfer may not be efficiently driven by ATG2A^NR^ alone. This appears consistent with previous *in vitro* reports of lower lipid transfer activity of *sp*Atg2^NR^ and ATG2A (AA 1-345) fragment than the full-length protein [20, 35] Collectively, these results suggest that regions other than the N-terminal domain are necessary for efficient phospholipid transfer.

Our MFEP also showed that the lipid prefers to pass through ATG2A^NR^ with the two acyl chains separated. At the entry step with low free energy barriers ∼8 kcal/mol, the phospholipid has to adjust its pose so that one acyl chain can insert deeper into the cavity. The subsequent flipping of the phospholipid into the cavity is the major bottleneck (∼15kcal/mol) for successful transfer because it requires expansion of the cavity for accommodating the lipid tail. Then the polar head has to bypass the protein helix, initiating curling of the lipid chain into the cavity, corresponding to a barrier of similar height. Finally, the phospholipid is completely engulfed by the cavity of ATG2A^NR^, reaching a reasonably stable transferred state **(Fig. 3c-d, video 3)**.

The detailed process of DOPC transfer by ATG2A^NR^ is as follows. Before DOPC enters the protein cavity, one of its acyl chains interacts with the protein. To initiate the process, this acyl chain has to find a suitable position through a series of fast conformational changes (free energy barriers <8 kcal/mol, **Fig. S6a-d**). Firstly, the polar DOPC head coordinates with both Arg18 and Trp86 **(Fig. S6a)**, which stabilizes its initial binding. Further, DOPC breaks its contact with Arg18, and then pushes Trp86 to rotate and leave space for its polar head to enter the protein cavity **(Fig. S6b-c)**. Next, the two acyl chains are separated, with the precursor chain entering the protein cavity and the polar head coordinated with Asp173 and Thr174 **(Fig. S6d)**. The next obstacle for DOPC is Phe171, which blocks the path of the polar head, creating a free energy barrier of ∼15 kcal/mol **(Fig. S6e)**. Then Phe171 flips outward to enlarge the internal space of the protein, allowing DOPC to detach from Thr174 for further insertion, and thus approaching Met125 **(Fig. S6f)**. Due to the absence of the downstream structure of ATG2A protein, the N-terminal cavity remains closed. Large hydrophobic amino acids, such as Trp121, blocks the entry of lipid molecules, resulting in a second high barrier of 15kcal/mol **(Fig. S6g)**. Eventually, after small scale rearrangement of DOPC in the cavity, Trp121 swings outward and the DOPC polar head forms a stable contact with Tyr200 on the β-sheet **(Fig. S6h)**. Detailed analysis of this process shows that the downstream protein integrity of ATG2A could be essential for efficient lipid transport. The presence of an intact cavity in ATG2A protein may effectively expand the internal space, and therefore decreases the energy barrier for lipid transfer.

To further consider the possible effect of the lipid environment in this process, we adopted the same TAPS approach but with two DOPCs **(Fig. S5d-f, video 4)**, one in the pocket of ATG2A^NR^ as before and the other staying at the transport entrance. Compared with the simulation of a single DOPC, the two DOPCs both stay at the N-terminus of the protein for much longer. In the former single-lipid setup, the only DOPC is in an aqueous environment before entering the protein cavity. Such aqueous environment seems more conducive to its movement into the hydrophobic pocket. The existence of a second lipid weakens such driving force, resulting in a high barrier at the initial step **(Fig. S7)**. In the first stable state, the carbon chains of the two molecules are entangled due to hydrophobic interactions and compete for entering **(Fig. S7a)**. Escaping from the entanglement, one DOPC (yellow in **Fig. S7b**) has to cross a high barrier so that its outer chain can bypass and push the DOPC (green) farther away from the protein to monopolize the entrance **(Fig. S7b)**. Subsequently, this precursor DOPC adjusts its pose and coordinates with Arg18 for stability **(Fig. S7c)**. After the precursor DOPC enters the protein, one of its hydrophobic chains occupies the entrance, indicating continuous lipid transport **(Fig. S7d)**. Except for the higher energy barrier in the beginning, the transfer process of the precursor DOPC closely resembles the case of single DOPC with barriers of similar height. This suggests that the retention of the N-terminal lipid environment has no obvious effect on the size of the protein cavity and the transfer of lipids after they enter ATG2A^NR^.

Note that, during lipid extract and transfer process, the polar headgroup of the lipid moves along the edge of the β-sheets, coordinated by a series of polar residues in the cavity, including Lys13, Arg15 during extraction and Tyr200/Lys255 during the transfer **(Fig. 3e)**. To verify these findings, we designed three mutants K13D, R15D and Y200E/K255E and examined their lipid transfer activities *in vitro* **(Fig. S8a)**. The highly curved small unilamellar vesicles (SUVs) meet the curvature requirement for phospholipid transfer by ATG2A, and were used as the lipid donor and acceptor. The SUVs were composed of 75% DOPC, 15% DOPE, 8% NBD-PE and 2% Rhodamine-PE for the donor, and 75% DOPC, 25% DOPE for the acceptor. The NBD-PE transfer from donor liposome to acceptor liposome by ATG2A results in the de-quenched NBD-florescence **(Fig. 3f)**. The wild type ATG2A can transferred NBD–PE from donor liposomes to acceptor liposomes. All three mutants showed substantially reduced activity relative to the wild type protein, which confirms the simulation predictions **(Fig. 3g).** Meanwhile, the acceptor liposome containing 75% DOPC, 20% DOPE and 5% PI3P was used to perform lipid transfer assay with similar results **(Fig. S8b-c)**.

### The interaction of ATG2A and WIPI4

WIPI4 is a β-propeller protein which binds to PI3P, and helps ATG2A anchoring to the phagophore membrane, thereby assisting in the phospholipid transfer process. The ATG2A-WIPI4 interaction has been studied previously [17, 41], however the molecular details remain unclear. In our ATG2A-WIPI4 density map, WIPI4 has much lower resolution than the rest of the protein **(Fig. 1c).** This is caused by lack of particle side views **(Fig. S2d)** but also due to the ability of WIPI4 to adopt a range of orientations relative to ATG2A, as observed in 2D class averages **(Fig. 4a)**. We then analyzed our XL-MS data for intermolecular cross-links, providing spatial restraints between pairs of amino acids located in ATG2A and WIPI4. Two inter-molecular crosslinks were identified between ATG2A K1203–WIPI4 K89 and ATG2A K1561–WIPI4 K40. WIPI4 K89 is located in blade 2, while K40 is located in blade 1. ATG2A residues K1203 and K1561 are located within 32 Å distance from WIPI4. With these new data providing experimental restraints, we rigid-body fitted a WIPI4 AlphaFold model in a density map representing one main conformation of ATG2A-WIPI4 complex. In the resulting model, blade 2 of WIPI4 interacts with the ATG2A β-sheet, while the PI3P binding blades 5 and 6 are at the opposite position **(Fig. 4b-c)**. This result is consistent with previous studies showing that ATG2A binding region is primarily located in blade 2 of Atg18 [42], and blades 1-3 of WIPI3 [41]. Interestingly, the main WIPI3/WIPI4 interaction site was previously narrowed down to the WIR-peptide of ATG2A (residues 1378 to 1399) [41]. This sequence is located in an extended loop that is not resolved in density maps. However, our data show that WIPI4 forms an additional interaction with several β-strands lining the C-terminal wall of the ATG2A lipid transfer cavity **(Fig. 4d)**. This contact appears to be a pivot point for WIPI4 movements relative to ATG2A **(Fig. 4a)**, allowing for close interaction between the two proteins and conformational adjustments at the membrane surface.

**Fig. 4:**
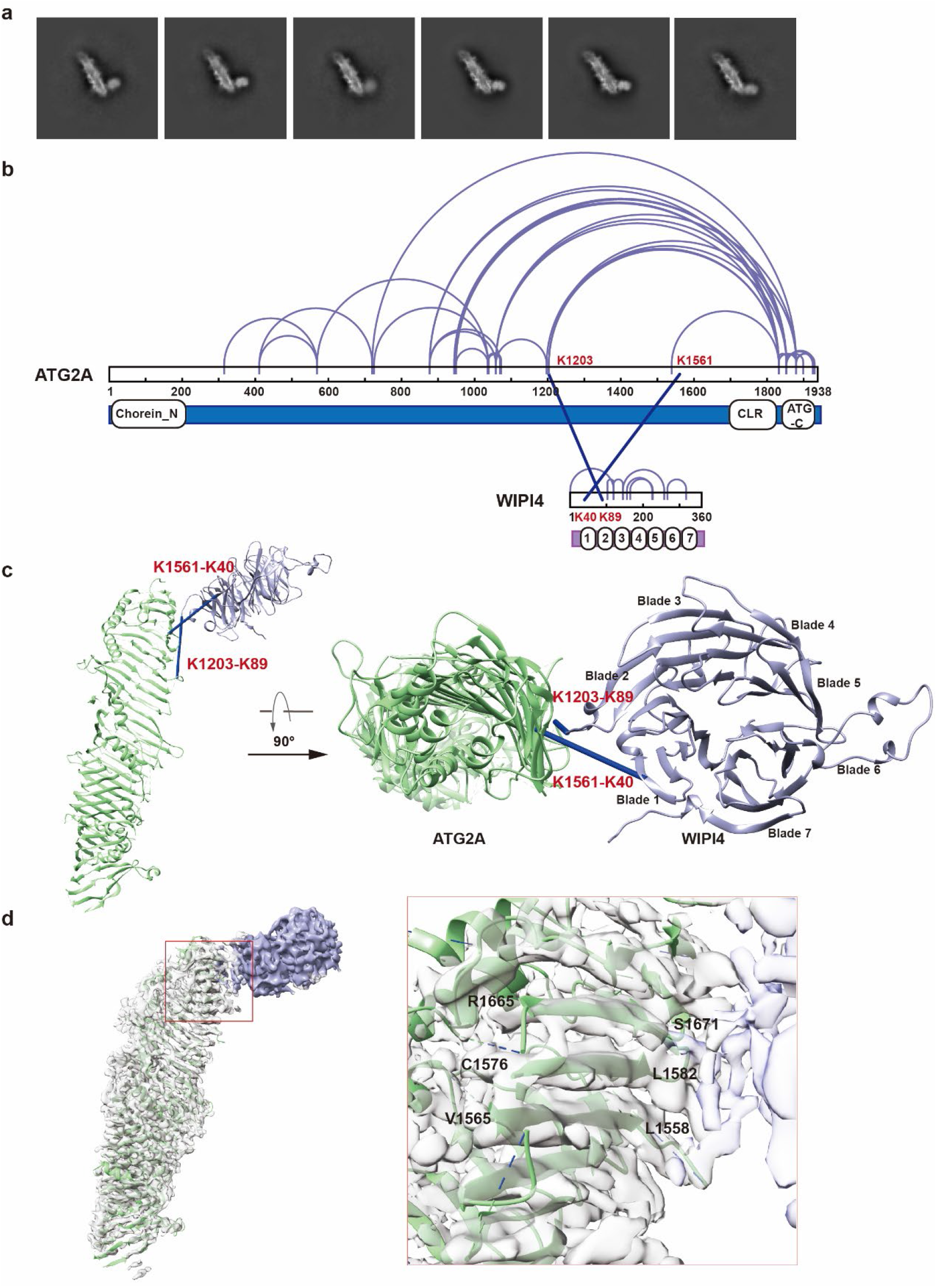
The interaction of ATG2A and WIPI4. **a,** Selected 2D class averages of the ATG2A-WIPI4 complex indicate the flexible binding of WIPI4. **b,** XL-MS analysis of ATG2A-WIPI4 complex. Linear plots displaying all the identified intramolecular (purple lines) and intermolecular (blue lines) crosslinks. **c,** Satisfied intermolecular XLs mapped onto the structures of ATG2A-WIPI4 complex. **d,** WIPI4 interaction interface is shown on the ATG2A model (green).

### Cryo-electron tomography reveals several membrane engagement modes of ATG2A-WIPI4

The cryo-EM structure of the ATG2A-WIPI4 complex revealed that the membrane-binding domains were flexible with respect to the central part of ATG2A, highlighting the possibility of flexibility in how the entire complex engages membranes. To test this, we performed cryo-ET on SUVs mixed with purified ATG2A-WIPI4. The cryo-electron tomograms clearly resolved the ∼20 nm diameter SUVs, as well as densities whose length closely corresponded to single copies of the ATG2A-WIPI4 complex at 20 nm **(Fig. 5a, Fig. S10a)**. Consistent with the robust lipid transfer activity in the assay, the tomograms revealed abundant examples of vesicle-vesicle tethering by ATG2A-WIPI4 **(Fig. 5a-b)**. Both the rod-shaped ATG2A and the WIPI4 density bound to its C-terminal side were clearly seen in several places in the tomograms **(Fig. 5a-b)**. This allowed analysis of the complex’s binding mode to vesicles. Strikingly, ATG2A-WIPI4 was observed to bind at several different angles to the tethered SUV membranes **(Fig. 5a-b, Fig. S10b-c)**. Fitting the ATG2A-WIPI4 model into the tether densities in the tomograms, we identified the conventional upright bridge-like model of the lipid transfer, where ATG2A is aligned orthogonally to the membranes, tethering them through the interaction of WIPI4 and its C-terminal domain to one SUV membrane and with its N-terminal binding to the opposite membrane **(Fig. 5c)**. In addition, ATG2A-WIPI4 complexes were found to lie parallel to the tethered membranes **(Fig. 5d)**. In this conformation the SUVs were brought in close proximity, separated by approximately 10 nm. An analysis of 100 tethering events in the tomograms revealed that ATG2A is found at a continuum of angles with respect to the membrane, with near-orthogonal binding to the membrane being the most prevalent orientation **(Fig. 5e)**. Taken together, cryo-ET analysis of ATG2A-WIPI4 interaction with SUVs revealed that ATG2A-WIPI4 can tether membranes in several different orientations, some or all of which may be active in lipid transfer.

**Fig. 5:**
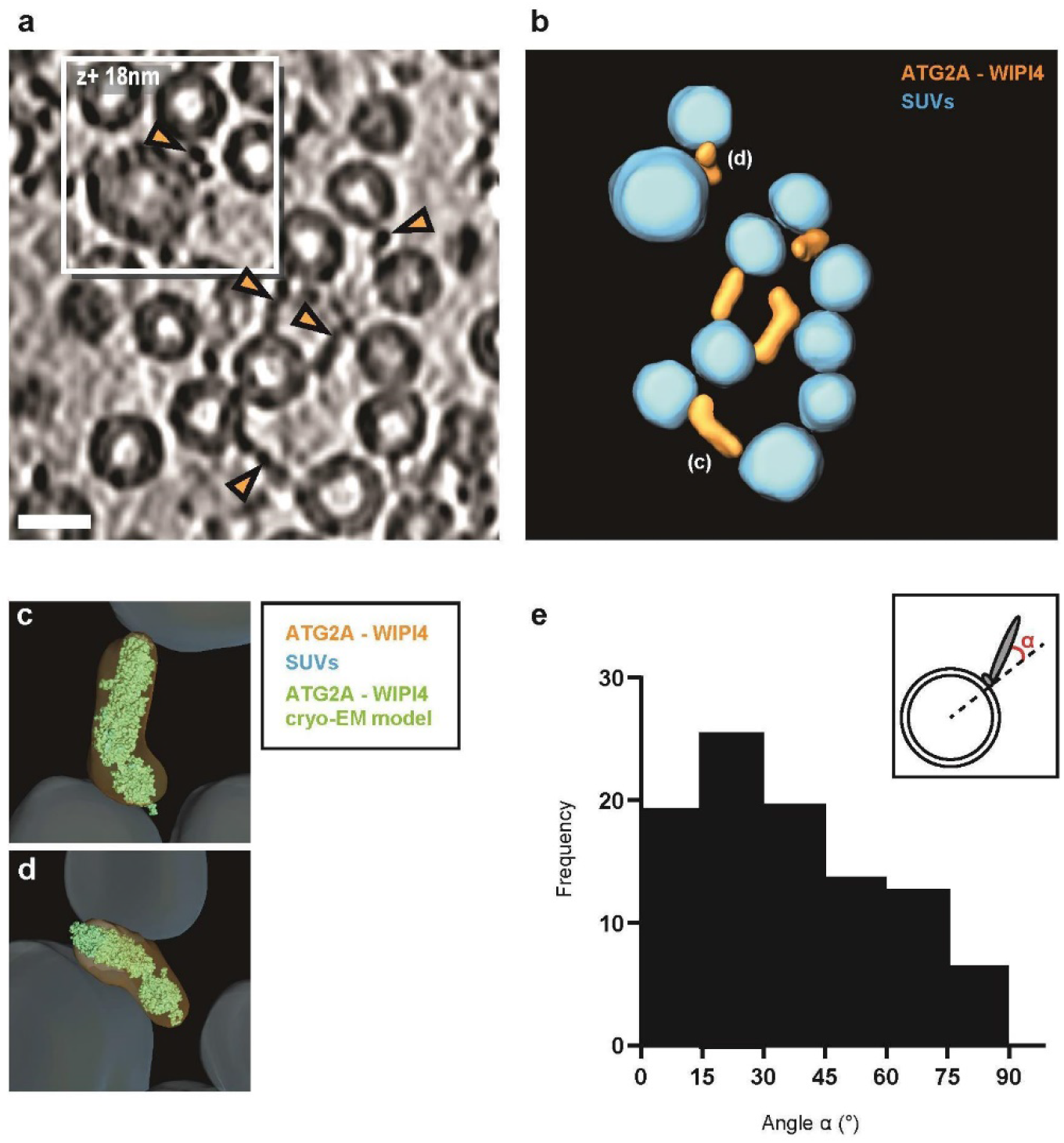
Cryo-electron tomography reveals several membrane engagement modes of ATG2A-WIPI4. **a,** Slice through a representative cryo-electron tomogram of decorated SUVs with ATG2A-WIPI4 protein complex. Orange arrowheads indicate ATG2A-WIPI4 densities tethering the SUVs. Density is dark. The tomogram was denoised using IsoNet [93]. The unfiltered version is shown in Fig. S10D. Scale bar, 20 nm. **b,** 3D segmentation of the tomogram presented in **(a)**. Color labels are defined for each structure. **c-d,** Magnified segmentations of ATG2A-WIPI4 complexes bound perpendicular (**c**) and flat (**d**) to SUV membranes, where the cryo-EM model of ATG2A-WIPI4 complex was fitted to its corresponding segmented volume. **(e)** 100 ATG2-WIPI4 densities were selected from six tomograms, and their relative orientation to the vesicle membrane was evaluated as indicated in the inset. Note that 0° corresponds to a perpendicular binding mode and 90° to a binding along the membrane tangent.

### The architecture of the ATG2A-WIPI4-ATG9A complex

ATG9A is a multi-span transmembrane protein that functions as a lipid scramblase, responsible for phospholipid redistribution between the inner and outer leaflet of the expanding phagophore membrane [23, 24]. ATG9A is trafficking between Golgi derived vesicles (ATG9 vesicles), phagophore and other membranes that may serve as lipid sources for autophagosome expansion [43]. It has been reported that Atg9 establishes Atg2-dependent contact sites between ER and phagophore through direct interaction, and this interaction is essential for autophagy [18]. The direct interaction raises the possibility that phospholipid transfer by ATG2A may be spatially coupled to the scramblase activity of ATG9A. To unveil the structural details of ATG9A-ATG2A-WIPI4 complex, we reconstituted the complex *in vitro* and studied its structure by cryo-EM **(Fig. S11)**. 2D class averages of the complex show that one ATG9A trimer can only bind one ATG2A through a single subunit, and the other two subunits are less resolved **(Fig. S12a, S13a)**. The GABARAP binding site is located in an extended loop [14] that was not resolved in 2D class averages or 3D reconstructions, probably due to high flexibility. Similar to ATG2A-WIPI4 complex alone, the WIPI4 density was readily observable bound to ATG2A in 2D class averages of the full complex. Since WIPI4 binds to the C-terminal end of ATG2A we could determine the ATG2A orientation relative to ATG9A. These results showed that one subunit from the ATG9A trimer can interact with both the N-terminal and the C-terminal end of ATG2A **(Fig. S12a, S13a)**, and therefore possibly form even larger assemblies with one ATG2A molecule bridging two ATG9A trimers. Indeed, we could observe 2D class averages with ATG9A densities flanking both ends of ATG2A **(Fig. S13c)**. These results are consistent with previous studies showing that both the N-terminal region (AA 237–431) [44], and C-terminus/CLR domain of Atg2/ATG2A [18, 45] are required for interaction with ATG9A. To verify the N-terminal binding model, the interaction between full-length ATG2A and N-terminal ATG2A construct (mini ATG2A, AA 1-443) with ATG9A was tested by *in vitro* pulldown assay. The results show the mini ATG2A can bind to ATG9A in amounts comparable with the full-length protein **(Fig. S14)**. Next, we performed 3D single particle reconstruction of ATG2A-ATG9A complexes. The nominal resolution for C-terminally and N-terminally bound ATG2A-ATG9A complex is ∼7.0 Å and ∼7.1 Å, respectively. In the case of the N-terminally bound complex, the ATG9A trimer density was poorly resolved and may be more dynamic. However, the interaction interface in ATG9 appears to be similar in both reconstructions **(Fig. 6a, Fig. S12a, S13a, d)**. Experimentally derived ATG2A and ATG9A models were rigid-body fitted in the ATG2A-ATG9A complex map **(Fig. 6a, Fig. S13d)**. In our model, ATG2A makes direct contact with one protomer of ATG9A through the C-terminal opening of the lipid transfer cavity. ATG2A is positioned nearly parallel to the predicted membrane plane, and binds to cytoplasmic domain of the ATG9A protomer, proximal to the lateral pore **(Fig. 6b)**. This interaction results in close alignment of the putative lipid transfer cavity opening with the lateral pore of ATG9A, thought to be important for lipid binding and translocation **(Fig. 6c)**.

**Fig. 6:**
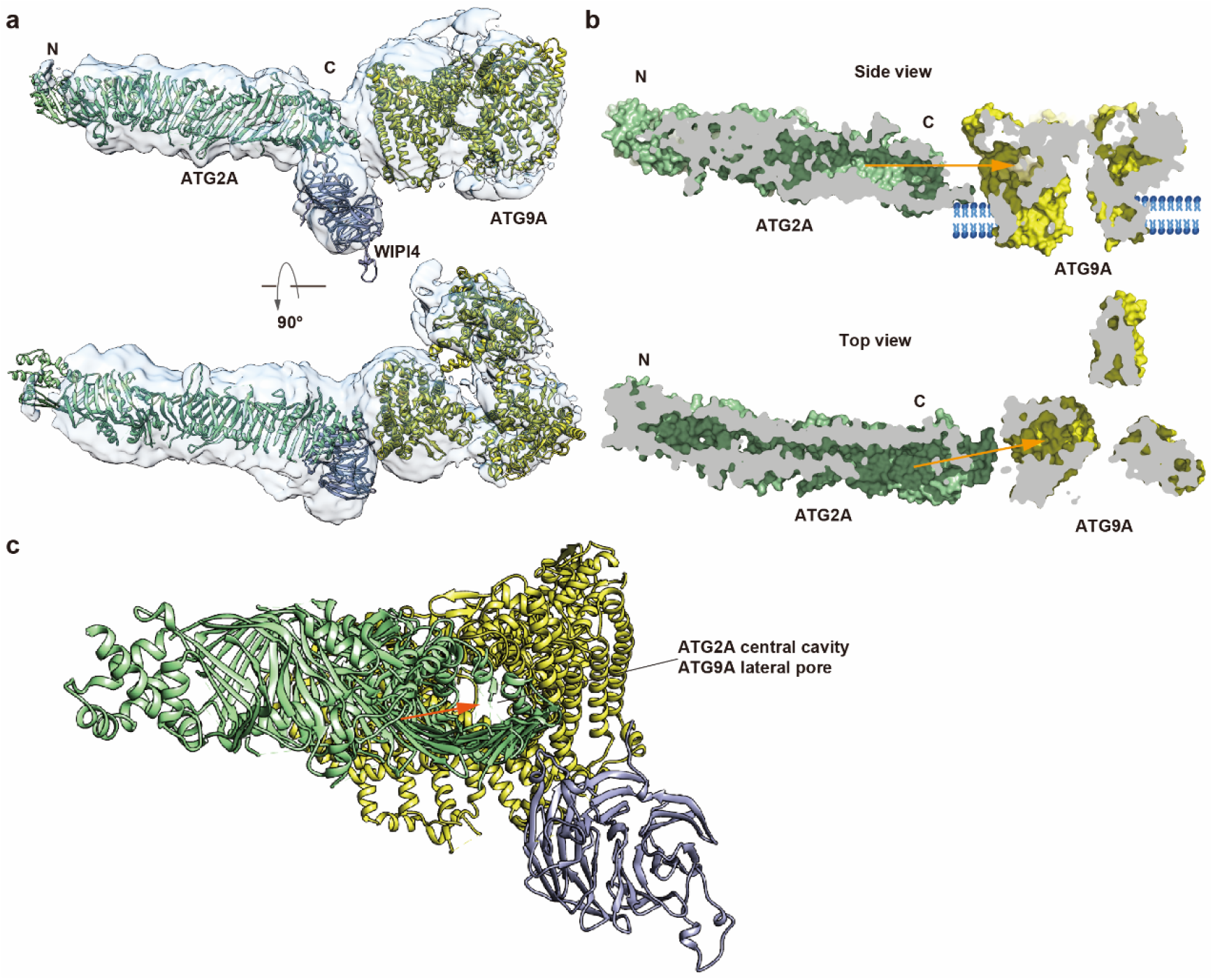
The cryo-EM structure of ATG2A-WIPI4-ATG9A complex. **a,** ATG2A C-terminal end binds in the close proximity of ATG9A lateral pore. ATG9A (yellow) and ATG2A-WIPI4 (green and purple) models were rigid body fitted in the electron density map. **b,** The cross-section of ATG2A-ATG9A surface is shown from side and top. **c,** Side view showing ATG2A central cavity alignment with the ATG9A lateral pore. ATG9A cytoplasmic domain binds to the C-terminus of ATG2A. ATG2A is colored in green, and ATG9A in yellow. Phospholipids exiting ATG2A hydrophobic cavity may directly enter into ATG9A lateral pore and redistribute to phagophore inner and outer leaflet.

Two groups have reported cryo-EM structure for human ATG9A homotrimer that confirmed broad features of the ATG9A structure reported here **(Fig. S15a)**. It is reported the open and closed conformation of ATG9 with 8° tilting of each protomer embedded in Lauryl Maltose Neopentyl Glycol (LMNG) [23]. Maeda et al. resolved three ATG9A timer structures purified with LMNG, amphipols and nano-discs [24]. Their amphipol and nano-disc structures are nearly identical in closed state, and LMNG structure assumes the open state [24]. We compared our structure of ATG9A embedded in LMNG with Maeda et al.’s open state ATG9A model (7JLQ). Results shows that our ATG9A model is in a more open state than previously reported. By aligning with each protomer, the movement between the neighboring monomer was 6.5 Å, 15.8 Å and 17.0 Å **(Fig. S15b)**. This movement results in larger displacement of one of the protomers in relation to the remining two, leading to the central pore radius widening **(Fig. 7e)**. To determine the ATG9A state in the C-terminally binding mode, ATG2A-WIPI4 and ATG9A map were refined separately by particle subtraction and local or non-uniform refinement **(Fig. 7a-b, Fig. S12a)**. Our open state ATG9A and previously reported close state ATG9A map (7JLP) were rigid-body fitted into the ATG2A-WIPI4-ATG9A map. In contrast to close state ATG9A map, the open state ATG9A and ATG2A-WIPI4-ATG9A map showed good overlap, with ATG9 central pore open in both cases **(Fig. 7c-d)**.

**Fig. 7:**
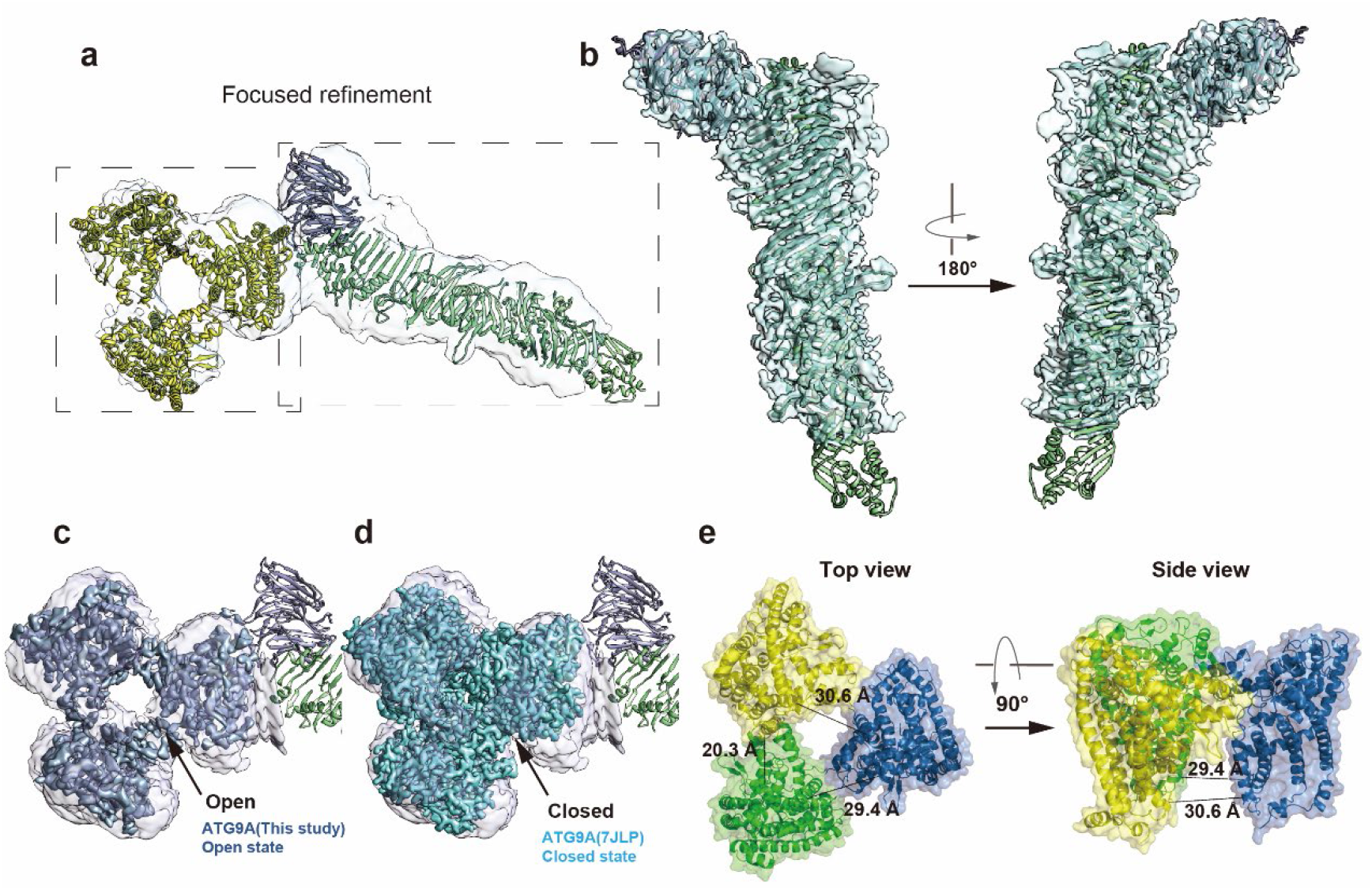
Open state of ATG9A bound to ATG2A/WIPI4 complex. **a,** The focused refinement of ATG2A-WIPI4 and ATG9A was performed separately. **b,** The local refined ATG2A-WIPI4 map with fitted model. **c,** The open state ATG9A map (blue) was superimposed with the focus refined ATG9A map (light purple) from the whole complex. **d,** The top view of closed state of ATG9A map fitted into focus refined ATG9A map. The closed state map was generated from reported model 7JLP by Chimera at the same resolution of open state ATG9A map. **e,** Distance between ATG9A protomers, which are colored in yellow, green and sky-blue. The distance was measured between I396 and T409 in each promoter.

Each of the ATG9A monomers contains one lipid binding pore, that opens laterally to the cytoplasmic leaflet. These three lateral pores are connected to a single central pore formed by physical interaction of all three monomers, at the trimer center, traversing both leaflets of the membrane. The close alignment of the ATG2A lipid transfer cavity and the ATG9 lateral pore observed in the 3D reconstruction would allow for phospholipids to transfer through ATG2A and then directly enter into ATG9A lateral pore and redistribute to phagophore inner and outer leaflet **(Fig. 6b-c)**. This spatially coupled mechanism may highly improve the lipid transfer efficiency. Furthermore, our results indicate that ATG2A N-terminus could also bind to ATG9A in the proximity of the lateral pore **(Fig. S13d)**, where ATG9A could serve as a lipid donor by directly feeding phospholipids from ATG9-positive vesicles into ATG2A to the phagophore membranes. This model is consistent with a recent study showing that Atg9 can facilitate lipid transfer when present in either donor vesicles, or in both donor and acceptor vesicles [46].

## Discussion

The source of phospholipoids for *de novo* autophagosome formation is a long-standing question in autophagy research. Recent studies have identified ATG2 as a key protein responsible for direct phospholipid transfer from ER membranes to the phagophore [17, 20, 35]. In addition to ER, Golgi, mitochondria, lipid droplets, and plasma membrane are also some of possible lipid sources for phagophore expansion [47]. ATG2A is a rod-like protein with a central hydrophobic cavity, proposed to bridge ER and autophagosome membranes via its N- and C-terminal ends, respectively [21]. The molecular mechanism of lipid transfer by ATG2A was hitherto limited to low-resolution EM structures [17, 20, 21]. Our ATG2A cryo-EM structure reveals how the extended network of β-strands folds into twisted β-sheets, forming a ∼200 Å long hydrophobic groove suitable for binding acyl chains and transferring multiple phospholipids to the acceptor membrane **(Fig. 1b, 2a)**. The groove side opening is solvent exposed and together with polar residues well suited to stabilize phospholipid head groups. This model may facilitate transferring process at a high rate of hundreds lipids per second as previously reported [48]. A conserved N-terminal domain of yeast Atg2 binds directly to the ER, and mediates lipid extraction from the membranes [35]. MD simulations and the FRET-based lipid transfer assay indicate that ATG2A^NR^ is similarly able to extract and initiate lipid transfer between the membranes. This process occurs in several steps, starting with acyl chain extraction by the N-terminal helical region, followed by phospholipid entry, flip, and finally transfer to hydrophobic groove of ATG2A **(Fig. 3d, video 3)**. This is consistent with recent studies on lipid binding and transport by ATG2A, showing that the first 345 residues are sufficient for lipid transport *in vitro* [20]. Rapid phagophore expansion requires unidirectional lipid transfer from the ER or other [34, 49]. The phagophore-bound receptors GABARAP and WIPI4, membrane lateral pressure and lipid biosynthesis may all contribute to unidirectional lipid transport. WIPI4 adopts a range of orientations, while maintaining close interaction with the C-terminal end of ATG2A rod-like domain **(Fig. 4a, 4d)**. This may allow ATG2A to dynamically associate with donor membranes while staying stably anchored on expanding phagophores or ATG9A-vesicles forming the seed membrane during the early stages of autophagosome formation. This is supported by cryo-ET analysis showing that ATG2A-WIPI4 can tether membranes in several different orientations **(Fig. 5)**. Our structural analysis provides the molecular details of the ATG2A-ATG9A complex and spatially coupled lipid transfer and scrambling activity. ATG2A interacts with one ATG9A monomer, while the other two ATG9 subunits appear to be in different conformations **(Fig. 6a, S12a)**. A recent study employing integrative modeling of the ATG2A-ATG9A complex showed a similar binding stoichiometry [45]. Our cryo-EM data suggest that ATG9 can interact with both the C-terminal and N-terminal parts of ATG2A **(Fig. S12, S13)**. This dual interaction mode is consistent with previous studies [18, 44, 45], and may play a role in unidirectional lipid transfer during initial and later stages of phagophore growth. ATG2A in 3D reconstruction is positioned nearly parallel to the predicted membrane plane **(Fig. 6b)**. We propose that this is only one of possible orientations that may be influenced by interactions with donor and acceptor membranes, as observed in our cryo-ET reconstructions of ATG2A-WIPI4 engagement with membranes **(Fig. 5)**. Therefore, the orientation of ATG2A relative to ATG9A probably changes in the native membrane environment due to the dynamic interaction interface. The lipid transfer by ATG2A is believed to be unidirectional in order to drive phagophore expansion from small ATG9-positive seed vesicles. The lipid synthesis, membrane curvature and the concentration of membrane proteins influence the membrane tension and may contribute towards unidirectional lipid flow through ATG2 [12, 50–52]. Lipid scramblases may also contribute to this process by equilibrating lipids between leaflets in both donor and acceptor membranes [32, 34, 50]. Future work is required to fully clarify the driving force behind unidirectional lipid flow through ATG2. Our data suggest that lipids are delivered to the lateral pore of ATG9A through direct interaction with ATG2A and close alignment with the lipid transfer cavity orifice. We propose that lipids are extracted from donor membranes such as ER by the ATG2A N-terminal domain, then transferred through the hydrophobic groove, and finally delivered to the ATG9A lateral pore **(Fig. 8a)**. Our model differs from a previously reported integrative modelling study that proposed delivery of lipids from ATG2A to the ATG9A perpendicular pore [45]. ATG9A lateral pores are directly connected to the central pore and allow for lipid flow and redistribution to phagophore inner and outer leaflet [23, 27]. This spatially coupled process can improve the lipid transfer efficiency [46, 53]. Furthermore, it was recently proposed that coupling of lipid transfer proteins and scramblases allows for the formation and expansion of phagophore double membrane while maintaining the volume of the enclosed content within the membrane relatively constant [32]. In addition to the seeding function, ATG9-positive vesicles also may serve as a membrane source for phagophore expansion [17, 29, 30, 44] or dynamically interact with phagophores without being incorporated into them [29]. ATG9A interaction with N-terminal domain of ATG2A may be functionally similar to the recently described interaction with VAMP1 and TMEM41B [25]. VAMP1 and TMEM41B are ER resident lipid scramblases that interact with the ATG2A N-terminal part, and are suggested to re-equilibrate the leaflets of the ER as lipids are extracted and transferred to ATG2A-ATG9A complex in acceptor membranes. Similarly, ATG9A may re-equilibrate the leaflets of the ATG9A-positive vesicles as lipids are extracted and transferred to the expanding phagophore edge by ATG2A and possibly have additional roles in the nucleation stage of the autophagosome formation **(Fig. 8b)**. The majority of ATG9 is present in a peripheral pool of ∼30-60 nm cytosolic vesicles [26, 30, 54], and our model implies reduction in the vesicles size as a consequence of lipid depletion as autophagosomes grow. ATG9 itself may by recycled through retrograde transport to the endosome or Golgi apparatus. Recent study showed that Atg2 lipid transfer efficiency increases when Atg9 is present in donor, in acceptor, or in both donor and acceptor vesicles [55]. Furthermore, the mitochondria-associated ER membrane, ATG2A N-terminus interact with ATG9A to promote ATG9A-vesicle delivery and regulate phagophore expansion [44]. The chorein family lipid transfer protein VPS13A has been reported to transfer lipids with the similar partnership. VPS13A can anchor to ER by N-terminal binding with VAP [56, 57], and locate to plasma membrane by C-terminal interacting with XK, a member of lipid scramblases family [58, 59]. Further studies will be required to determine how these lipid scramblases coordinate with each other and potentially drive unidirectional lipid transfer by ATG2A. Taken together, we provide new structural and mechanistic insights into ATG2A-ATG9A mediated lipid transfer and re-equilibration required for autophagosome formation.

**Fig. 8:**
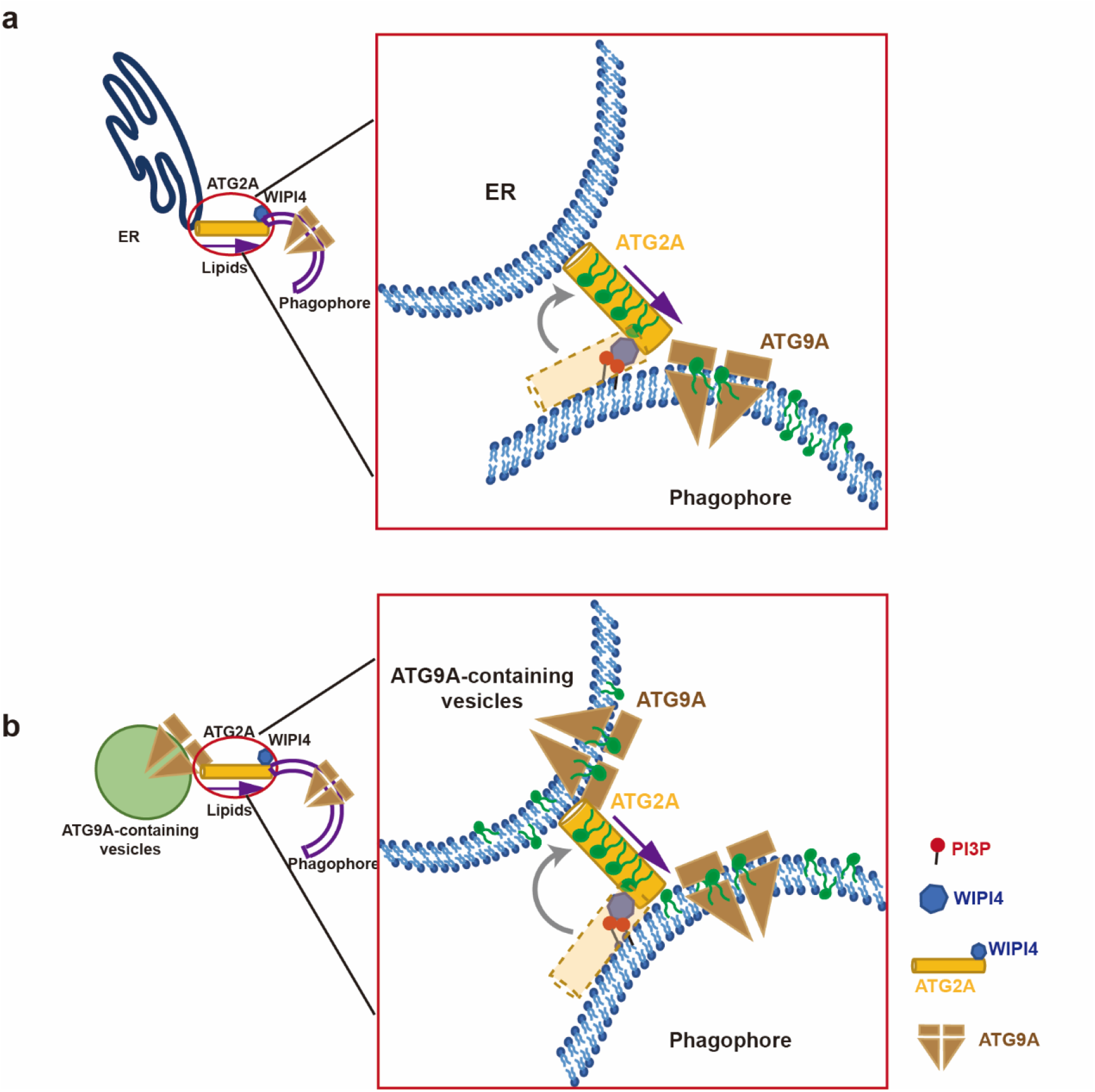
Proposed models of ATG2A-ATG9A mediated lipid transfer process. **a**, A proposed model of the lipid transfer machinery. N-terminal end of ATG2A interacts with ER or other organelles, while C-terminal regions interact with ATG9A and phagophore membrane. ATG2A can dynamically reposition in response to the native membrane environment. **b**, A proposed model of lipids transferred from ATG9A-containing vesicle to phagophore. The interaction of ATG2A and ATG9A promotes lipids transfer and re-equilibration between ATG9A-positive vesicles and expanding phagophore or another ATG9A-vesicle comprising the seed membrane.

**Table 1.**
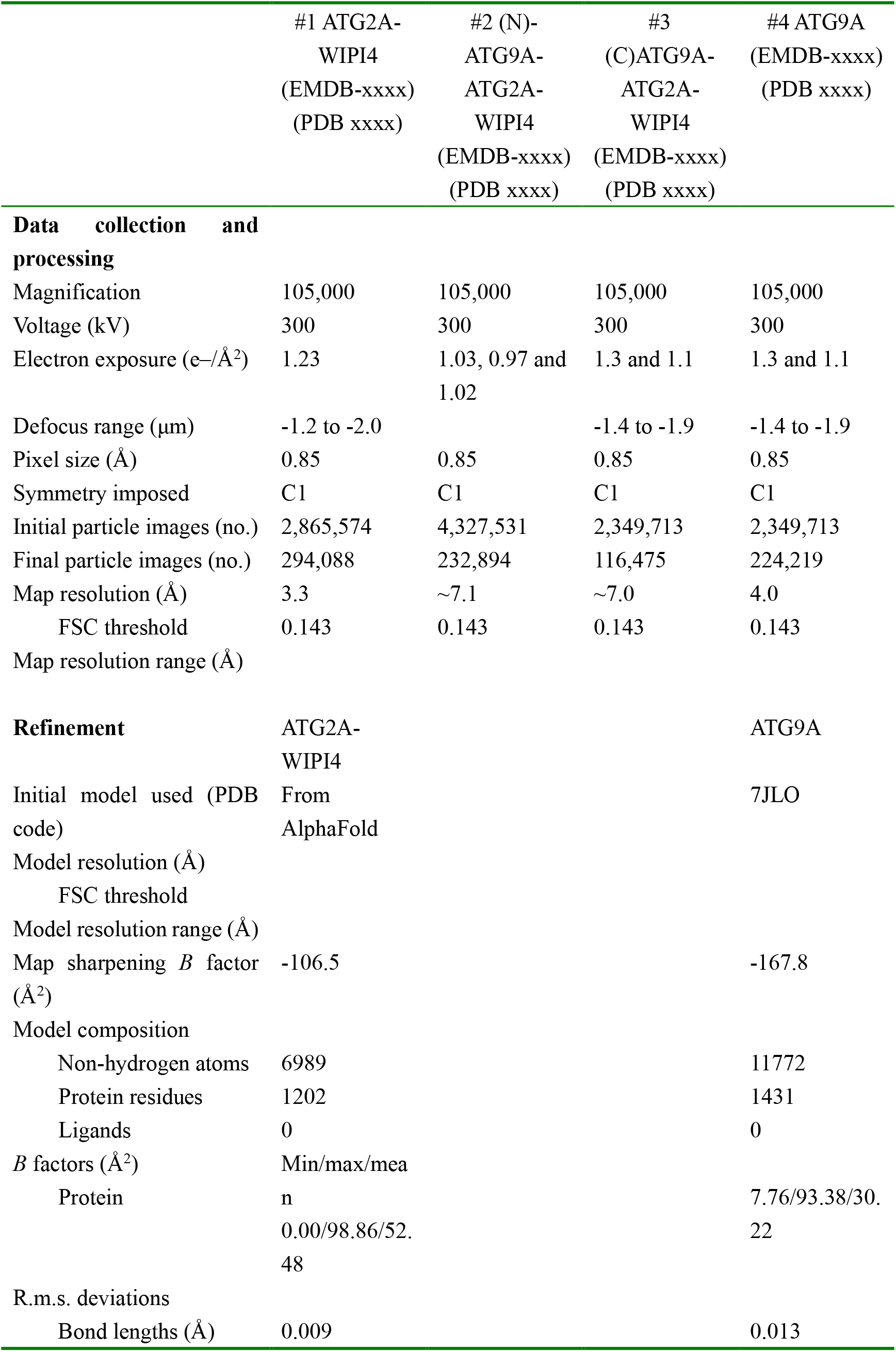
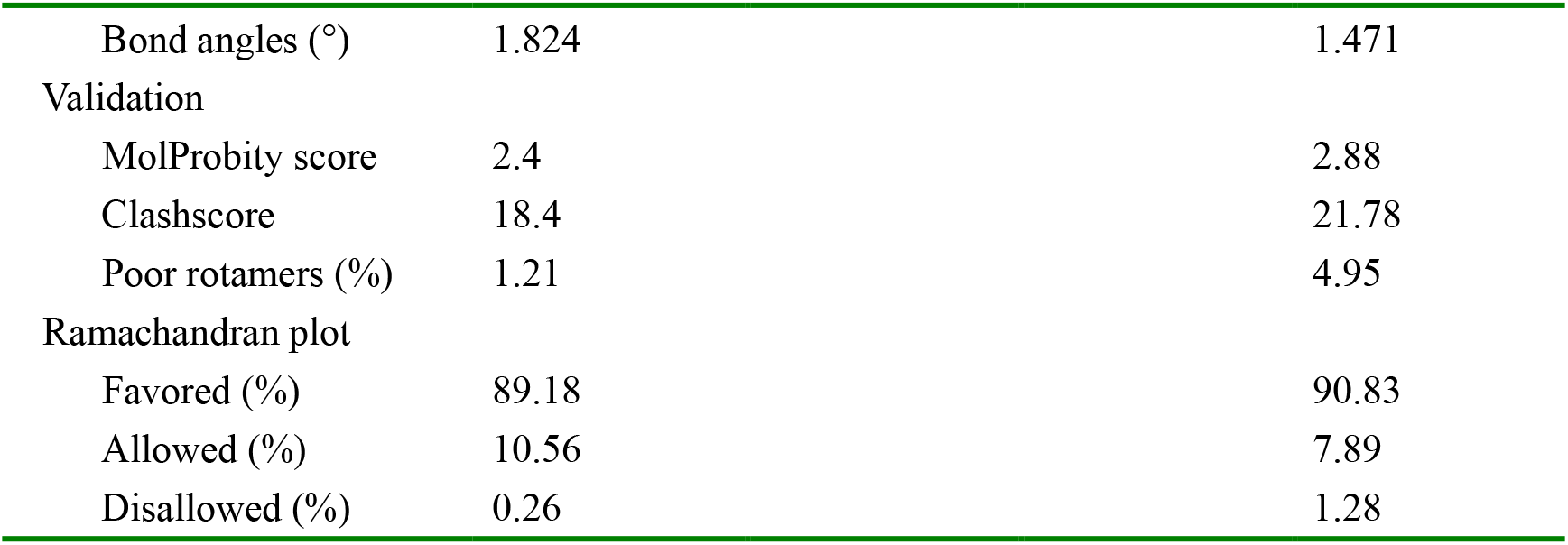
Cryo-EM data collection, refinement and validation statistics.

|  | #1 ATG2A-WIPI4<br>(EMDB-xxxx)<br>(PDB xxxx) | #2 (N)-<br>ATG9A-ATG2A-WIPI4<br>(EMDB-xxxx)<br>(PDB xxxx) | #3<br>(C)ATG9A-ATG2A-WIPI4<br>(EMDB-xxxx)<br>(PDB xxxx) | #4 ATG9A<br>(EMDB-xxxx)<br>(PDB xxxx) |
| --- | --- | --- | --- | --- |
| **Data collection and processing** |  |  |  |  |
| Magnification | 105,000 | 105,000 | 105,000 | 105,000 |
| Voltage (kV) | 300 | 300 | 300 | 300 |
| Electron exposure (e-/Å<sup>2</sup>) | 1.23 | 1.03, 0.97 and<br>1.02 | 1.3 and 1.1 | 1.3 and 1.1 |
| Defocus range (µm) | -1.2 to -2.0 |  | -1.4 to -1.9 | -1.4 to -1.9 |
| Pixel size (Å) | 0.85 | 0.85 | 0.85 | 0.85 |
| Symmetry imposed | C1 | C1 | C1 | C1 |
| Initial particle images (no.) | 2,865,574 | 4,327,531 | 2,349,713 | 2,349,713 |
| Final particle images (no.) | 294,088 | 232,894 | 116,475 | 224,219 |
| Map resolution (Å) | 3.3 | ~7.1 | ~7.0 | 4.0 |
| FSC threshold | 0.143 | 0.143 | 0.143 | 0.143 |
| Map resolution range (Å) |  |  |  |  |
| **Refinement** | ATG2A-WIPI4 |  |  | ATG9A |
| Initial model used (PDB code) | From<br>AlphaFold |  |  | 7JLO |
| Model resolution (Å) |  |  |  |  |
| FSC threshold |  |  |  |  |
| Model resolution range (Å) |  |  |  |  |
| Map sharpening *B* factor (Å<sup>2</sup>) | -106.5 |  |  | -167.8 |
| Model composition |  |  |  |  |
| Non-hydrogen atoms | 6989 |  |  | 11772 |
| Protein residues | 1202 |  |  | 1431 |
| Ligands | 0 |  |  | 0 |
| *B* factors (Å<sup>2</sup>) | Min/max/mea |  |  |  |
| Protein | n<br>0.00/98.86/52.<br>48 |  |  | 7.76/93.38/30.<br>22 |
| R.m.s. deviations |  |  |  |  |
| Bond lengths (Å) | 0.009 |  |  | 0.013 |

|  |  |  |
| --- | --- | --- |
| Bond angles (°) | 1.824 | 1.471 |
| Validation |  |  |
| MolProbity score | 2.4 | 2.88 |
| Clashscore | 18.4 | 21.78 |
| Poor rotamers (%) | 1.21 | 4.95 |
| Ramachandran plot |  |  |
| Favored (%) | 89.18 | 90.83 |
| Allowed (%) | 10.56 | 7.89 |
| Disallowed (%) | 0.26 | 1.28 |

## Methods

### Cloning

ATG2A optimized gene was cloned into pCAG vector, WIPI4 optimized gene was cloned into pFastBac-Dual vector, ATG9A optimized gene was cloned into pCAG vector, GABARAP gene was cloned from cDNA and inserted into pET28a vector.

### Protein expression and purification

ATG2A wild type and mutants had TEV-cleavable glutathione-S-transferase (GST) and Twin-StrepII-Flag at N-terminal, WIPI4 was GST tagged at N-terminal, GABARAP was 6xHistidine (His) tagged at N-terminal, ATG9A was 6xHis and Twin-StrepII tagged at N-terminal. ATG2A and ATG9A was expressed in HEK Expi293F cells, GABARAP was expressed in *Escherichia coli* BL21(DE3). WIPI4 was expressed in insect cell system, the baculoviruses were generated from the resulting vector in *Spodoptera frugiperda* 9 (Sf9) cells, and infected Hi5 insect cells.

For WIPI4 purification, the pellet was resuspended in lysis buffer 20 mM HEPES pH7.5, 150 mM NaCl, 5 mM DTT, 2% (v/v) Triton X-100 and 1 mM PMSF, 1 μM pepstatin, 10 μM leupeptin, 1 μg/mL aprotinin, followed by 1 hour stir-mixing at 4°C. The lysate was clarified by centrifugation at 18,000 rpm, and the protein in the supernatant was diluted to 1% Triton and incubate with Glutathione Sepharose beads (GE Healthcare) in a batch mode. After 2 hours of incubation at 4°C, the mixture was transferred to gravity column and then washed by wash buffer containing 20 mM HEPES pH7.5, 150 mM NaCl, 5 mM DTT. WIPI4 was eluted in Gel filtration buffer 20 mM HEPES pH7.5, 150 mM NaCl, 2 mM MgCl_2_, 1 mM TCEP and containing 50 mM glutathione, then TEV digested overnight. After buffer exchange to gel filtration buffer without glutathione by concentration, protein was incubated with GST resin again, further the collected flowthrough was injected into the Superose 75 Increase 10/300 column (GE Healthcare) **(Fig. S1b)**. The WIPI4 protein from gel filtration was flash-frozen in liquid nitrogen and stored at -80°C until further use.

ATG2A pellet was resuspended in lysis buffer 20 mM HEPES pH7.5, 500 mM NaCl, 5 mM DTT, 10% glycerol, 1% Triton X-100 and 1 mM PMSF, 1 μM pepstatin, 10 μM leupeptin, 1 μg/mL aprotinin. After centrifugation, the supernatant was incubated with GST beads for 2 hours at 4°C, later transferred to gravity column and washed with wash buffer containing 20 mM HEPES pH7.5, 500 mM NaCl, 10% glycerol, 5 mM DTT. After elution with gel filtration buffer containing 20 mM HEPES pH 7.5, 150 mM NaCl, 2 mM MgCl_2_, 1 mM TCEP supplemented with 50 mM glutathione, GST tag was digested by TEV overnight with 1 mM ATP to release chaperone. Next, the sample was loaded on Strep-Tactin Superflow beads (IBA), washed with gel filtration buffer with and without 1 mM ATP, then eluted with gel filtration buffer supplemented with 5 mM desthiobiotin. Finally, the protein was purified with Superose 6 Increase 10/300 column (GE Healthcare) **(Fig. S1a)**. For the ATG2A-WIPI4 complex purification, the purified WIPI4 was bound to ATG2A on Strep column, the overloaded WIPI4 was washed out before loading on Superose 6 Increase column **(Fig. S1c)**.

GABARAP pellet was resuspended by lysis buffer with 10 mM Tris, pH 7.0, 200 mM NaCl, 20 mM imidazole and 1 mM PMSF. The cells were sonicated at 200 W for 30 min, then centrifugated at 18,000 rpm for 40 min at 4°C. The clarified lysis was incubated with Talon (GE health) beads at 4°C and transferred to gravity column, washed with lysis buffer and eluted with elution buffer, 10 mM Tris, pH7.0, 200 mM NaCl, 200 mM imidazole. The protein was concentrated and purified on Superose 75 Increase 10/300 column with gel filtration buffer, 20 mM HEPES pH7.2, 200 mM NaCl, 2 mM MgCl_2_ **(Fig. S11a, d)**. ATG2A-WIPI4-GABARAP complex was reconstituted in a similar way as ATG2A-WIPI4 complex, purified WIPI4 and GABARAP were bound to ATG2A during Strep-Tactin affinity purification, then flash frozen and store at -80°C. ATG9A purification was referred to Guardia et al.’s work [23]. LMNG was used to extract protein from membrane and later purification. ATG9A was purified by Nickel affinity resin (QIAGEN) followed by Strep-Tactin and gel filtration with Superdex 200 10/300 column (GE Healthcare) **(Fig. S11b, e)**.

### ATG2A-ATG9A complex reconstitution

All purified proteins including ATG9A and ATG2A-WIPI4-GABARAP were flash frozen and stored at -80°C until use. The whole complex was reconstituted by pulldown with His beads. ATG9A and ATG2A-WIPI4-GABARAP (molar ration=1:1.5 and 1:3, ATG9A molecular weight calculated as a trimer) were incubated at 4°C for 2 hours with pre-equilibrated Nickel beads, followed by washing three times with 20 mM HEPES, pH 8.0, 150 mM NaCl, 1 mM TCEP, 0.09 mM LMNG, and eluting with 20 mM HEPES, pH 8.0, 150 mM NaCl, 200 mM imidazole, 1 mM TCEP, 1 mM EDTA, 0.09 mM LMNG **(Fig. S11c)**.

### Cryo-EM sample preparation and data collection

The freshly purified protein was used to freeze grids immediately, 3 μL of 0.3 mg/mL ATG2A-WIPI4 was applied to Quantifoil UltraAuFoil R1.2/1.3 300 mesh grids after glow discharge. Grids were plunged into liquid ethane using FEI Vitrobot Mark IV (Thermo Fisher Scientific) after blotting for 3 seconds at a blot force of 0 in 100% humidity at 4°C. A total of 6179 movies were collected on FEI Titan Krios G3 300-kV transmission electron microscope with K3 detection camera at a defocus range of -1.2 to -2.0 μm in counting mode. The magnification was 105kx leading to a specimen pixel size of 0.85 Å, and total exposure time was 2.5 seconds fractionated into 50 frames for a total dose of 61.65 e^−^ / Å^2^.

The freshly reconstituted ATG9A-ATG2A-WIPI4-GABARAP complex was diluted to 0.3 mg/mL and applied to Quantifoil UltraAuFoil R1.2/1.3 300 mesh grids after glow discharge. The vitrification and data collection conditions are the same as described above. For C-terminal binding ATG9A (ATG9A trimer: ATG2A complex molar ration=1:1.5), two sets of movies were collected at a total dose of 65.17 e^−^ / Å^2^ and 54.88 e^−^ / Å^2^ (50 frames), 4356 and 5856 movies separately, total 10212 movies. The defocus range was -1.4 to -1.9 μm. The N-terminal binding ATG9A group collected three datasets (molar ration=1:3), 7749, 6471 and 7047 movies, total 21267 movies. The total dose was 51.46, 48.39 and 51.1 e^−^ / Å^2^, the defocus was -1.2 to -1.8 μm.

### Cryo-EM data processing and 3D reconstruction

The data was processed by Relion 3.1.0 and cryoSPARC 3.2.0 program [60–63]. The movies were divided into five-by-five patches for motion correction by Relion’s own implementation of the MotionCor2 algorithm [62]. After motion correction, micrographs were transferred to cryoSPARC to proceed with contrast transfer function estimation by Patch CTF. The blob picker and template picker were applied to pick the particles and subjected to 2D classification.

For ATG2A-WIPI4 complex, 294,088 particles were selected from the best 2D class averages and put into Ab initial model and non-uniform refinement, which generated a 3.23 Å map. For ATG2A-WIPI4-GABARAP-ATG9A (C-terminus) complex was separated by 2D classification from the solo ATG9A and ATG2A-WIPI4. Then 116,475 particles were selected and applied to *ab initio* reconstitution and further refined by non-uniform refinement, which generated a map of 7 Å **(Fig. S12a)**. Meanwhile, for ATG2A-WIPI4-GABARAP-ATG9A (N-terminus) analysis, 232,894 particles were selected and reconstituted, and generated a map of 7.1 Å in CryoSPARC **(Fig. S13)**. For both N-terminus and C-terminus complex, the mask of ATG2A-WIPI4 and ATG9A were made separately, and performed local refinement and non-uniform refinement. Also, the particle subtraction was used to subtract ATG9A and ATG2A-WIPI4 individually, and further refined by local and non-uniform refinement. The refined map allows us to fit ATG2A-WIPI4 model with right handedness **(Fig. 7b)**.

The dissociated ATG9A subset from ATG2A-WIPI4-GABARAP-ATG9A (C-terminus) dataset was processed separately. Similarly, after 2D classification, ab-initial model and non-uniform refinement were performed, and a 3.97 Å resolution map was generated from 224,219 particles **(Fig. S12a)**.

### Model building and refinement

The initial model of ATG2A was based on the model in AlphaFold DB [64, 65]. Further model building into the cryo-EM map were performed by Coot and UCSF Chimera. The N-terminal region of ATG2A (8-236AA) model was homology-modeled by SWISS-MODEL [66]. PHENIX [67] and Coot [68] was used to do the real space refinement and manual modification, molecular graphics and analyses performed with PyMOL2.3.3 (The PyMOL Molecular Graphics System, Version 2.3.3 Schrödinger, LLC), UCSF Chimera1.14 [69] and UCSF Chimera X1.3 [70]. Due to the map resolution limitation, WIPI4 was rigid body fitted into the density map. For ATG9A-ATG2A-WIPI4-GABARAP complex structure, the experimental ATG2A-WIPI4 and ATG9A model was rigid body fitted into the density map by using global search fitting in Chimera [69]. The cryo-EM density maps have been deposited in the Electron Microscopy Data Bank under accession code XXXXX and the coordinates have been deposited in the Protein Data Bank under accession number XXXX.

### Molecular dynamics simulation

DOPC micelle was built by the Membrane Builder module in CHARMM-GUI server [71]. All MD simulations were performed using GROMACS-2019.4 [72]. The protein and lipids were solvated in a cubic box with TIP3P waters and 0.15 M Na^+^/Cl^-^ ions. The CHARMM36 force-field [73] was used to describe the interactions in the system. Energy minimization was performed for 10000 steps by the steepest descent algorithm and then by the conjugate gradient algorithm. Then a 100 ps NVT simulation was performed at 310 K for solvent equilibration, followed by a 1 ns NPT equilibration to 1 atm using the Berendsen barostat [74]. The production MD simulations were performed for 100 ns with a time-step of 2 fs and Parrinello-Rahman barostat [75]. In all constant temperature simulations, the Bussi (velocity-rescaling) thermostat was used [76]. Long-range electrostatic interactions were treated by the particle-mesh Ewald method [77, 78]. The short-range electrostatic and van der Waals interactions both used a cutoff of 10 Å. All bonds were constrained by the LINCS algorithm [79].

### Path searching

The initial and target structures were first equilibrated by short MD simulations. An initial transferring path was generated by targeted MD simulation via GROMACS-2019.4 and PLUMED-2.5.3 [80], in which the DOPC molecule was pulled from the N-terminal side to the hydrophobic cavity of ATG2A^NR^. The kappa value was set to 10000 kJ/(mol· *nm*^2^). The biasing force was applied to all C-alpha atoms of ATG2A^NR^ and all heavy atoms of DOPC, using the former for structure alignment.

The MFEP was located by the Traveling-salesman based Automated Path Searching (TAPS) algorithm [36]. The TAPS simulation was performed using an in-house python script incorporating GROMACS, PLUMED and Concorde [81] in the NVT ensemble at 310 K. The input path was obtained by selecting conformations from the targeted MD trajectory such that the root mean-square distance (RMSD) between neighbor conformations are 1.5Å. For TAPS, RMSD between any pair of conformations was computed using all heavy atoms of DOPC and inner side chains of ATG2A^NR^ **(Extended Table 2)** while structure alignment was performed using C-alpha atoms of ATG2A^NR^. In each TAPS iteration, 1000 ps sampling was performed in total. Gaussians of height 0.25 kJ/mol and width 0.5 were deposited on PCV_s [82] every 0.01 ps, with frames recorded at the same frequency. Convergence of TAPS was evaluated by Multidimensional Scaling (MDS) and the z component of the Path Collective Variable (PCV) [82].

### Free energy calculation

Umbrella sampling [40] was performed along the s component of the PCV of the MFEP found by TAPS via PLUMED, with wall potential limiting the range of sampling within 1.5Å from the MFEP (measured by √PCV_z). The RMSD definition and structure alignment remained the same as the TAPS optimization. The window size was chosen as 0.5 along the PCV_s of the MFEP. For each window, the strength of restrain was set as 100 kJ/mol. Each window was simulated for at least 10 ns. The free energy profile was obtained with the weighted histogram analysis method (WHAM) [83].

### Liposome preparation for lipids transfer assay

Small unilamellar vesicles were composed of phospholipids purchased from Avanti Polar Lipids. The donor liposomes were made up of 75% 1,2-dioleoyl-sn-glycero-3-phosphocholine (DOPC), 15% DOPE, 8% NBD–PE, and 2% Liss Rhod–PE, the acceptor liposomes were made up of 75% DOPC and 25% DOPE. The chloroform-dissolved lipids were mixed in a glass vial and dried under nitrogen gas stream, then dried further overnight under vacuum centrifuge. The lipid film was hydrated with ATG2A gel filtration buffer and the final suspension concentration was 0.5 mM. The lipids were resuspended at room temperature and vortexed for 2 min, followed by 10 times freeze-thaw treatment then sonicated at 100W, 2s on, 5s off for 40 cycles. Similar, the acceptor liposomes containing of 75% DOPC, 20% DOPE and 5% PI3P were also made for further activity assay.

### Lipids transfer activity assay

The lipids transfer reaction was prepared in 96-well microplate, the mixture was composed of 20 μM donor liposomes, 100 μM acceptor liposomes and 50 nM purified ATG2A variants. After addition of the proteins to the liposomes, the microplate was immediately set on fluorescence intensity reader (Perkin Elmer Multimode Plate Reader, EnVision 2105), monitored for 30min. Excitation wavelength was set at 460 nm; fluorescence emission was recorded at 535 nm. Next, NBD fluorescence intensity was quantified after 40 min incubation with 0.4% n-dodecyl-β-d-maltoside (DDM). For the activity assay, both ATG2A wild type and mutants were purified by GST and Strep-Tactin resin. The experiments were repeated three times, and shown as the mean value with standard error of the mean (SEM).

### Crosslinking mass spectrometry (XL MS)

16 μg of ATG2A-WIPI4 at 0.5 μM concentration was crosslinked by 0.2 mM DSBU (Thermo Scientific) at RT for 40 min **(Fig. S9)**. After reaction, 1 M Tris was used to quench the reaction on ice for 30min with final 100 mM concentration. Crosslinked protein was dried in vacuum centrifuge for 3 hours and redissolved in 100 μL 8M urea, 50 mM Tris-HCl (pH 8.0), 10 mM DTT, mixed briefly and incubated at 37°C for 60 min. When the sample cooled to RT, chloroacetamide (CAA) was added from the 500 mM stock solution to a final concentration of 50 mM, mixed and incubated at RT in the dark for 30 min. For trypsin digestion, sample was diluted with 50 mM Tris-HCl (pH 8.0), 1 mM CaCl2 solution to a final concentration of 1 M urea, and then added trypsin at an enzyme-to-substrate ratio of 1:20 (w/w). Mixed briefly by vertexing, and incubated the mixture overnight at 37°C. The digested sample was desalted using Pierce peptide desalting spin columns, separated by EASY nLC 1200 and analyzed by Orbitrap Eclipse mass spectrometer (Thermo Fisher) using sceHCD-MS2 fragmentation approach. Survey scans were recorded in the Orbitrap at 60 000 resolution (AGC 4e5, max injection time 50 ms) and a scan range from 350 to 1500 m/z. MS/MS scans were recorded in the Orbitrap at 30 000 resolutions (AGC 1e5, max injection time 200 ms, isolation width 1.6 m/z). Unknown, singly and doubly charged ions were excluded from fragmentation. Selected precursors were fragmented by applying a stepped-HCD energy of 30 ± 3% NCE. Proteome Discoverer 2.5 (Thermo Fisher) with XlinkX software 2.5 was used to analyze and identify the crosslinking peptides and the parameters were as follows: maximum of three missed cleavage sites for trypsin per peptide; cysteine carbamidomethylation as fixed modification and methionine oxidation as dynamic modification. Searches were performed against an ad-hock database containing the sequences of ATG2A-WIPI4 complex and common contaminant proteins (CRAPome/Strep tag AP [84]). Search results were filtered by requiring precursor tolerance (±10 ppm) and fragment tolerance (±20 ppm). FDR threshold was set to 1% at Crosslink and CSM level. Xlink Analyzer v1.12 as a plugin to UCSF Chimera was used to visualize the cross-links in model [85]. The mass spectrometry proteomics data have been deposited to the ProteomeXchange Consortium via the PRIDE [86] partner repository with the dataset identifier PXDXXXXXX.

### Sample preparation for cryo-electron tomography

SUVs for cryo-electron tomography were prepared from lipids purchased from Avanti Polar Lipids. The lipids were combined in a glass vial at a molar ratio of: 75% POPC, 20% POPS (dissolved in chloroform), 5% PI(3)P (dissolved in methanol). The lipid mixture was dried under a stream of nitrogen gas, and further dried under vacuum overnight. The lipid film was then hydrated with buffer (50 mM Tris-HCl pH 8, 150 mM NaCl and 0.1 mM TCEP) to a final lipid concentration of 12 mg/mL, thoroughly mixed, and heated to 45°C for 15 min. SUVs were formed using probe tip sonication for 15 min while cooled by an ice-water bath. The resulting SUVs were subjected to centrifugation 15,000 rpm for 15 min to pellet big vesicles. The supernatant containing SUVs (20 - 50 nm) were stored at 4°C.

For grid preparation, SUVs were mixed with ATG2-WIPI4 to a final concentration of 6 mg/mL SUVs and 2 µM ATG2A-WIPI4, and incubated on ice for 15 min. 10 nm protein A-coupled colloidal gold (CMC-Utrecht) was added to the final volume of 4 µL, applied to glow-discharged R1.2/1.3 Au (200 mesh) EM grids (QUANTIFOIL GmbH), after which the grids were immediately plunge frozen in propane-ethane using a FEI Vitrobot (FEI, ThermoFisher) at 22°C, 80% humidity, with blot force - 25 and a blot time of 2 seconds.

### Cryo-electron tomography data acquisition and processing

Data acquisition was performed on a Titan Krios electron microscope (ThermoFisher Scientific) operated at 300 kV using a GIF Bio-Quantum Energy Filter and K2 Summit detector (Gatan). 16 tilt series were acquired using SerialEM [87] in dose-symmetric mode [88], at a nominal magnification of 64,000 corresponding to a specimen pixel size of 1.1 Å in super-resolution mode, with spot size 9 and an objective aperture of 100 μm. The tilt range was set to ± 60° in 3° increments. The defocus setting was in the range of -3 to -4 μm. The total dose for a tilt series was 100 e^−^/Å^2^. Movie stacks were drift corrected using MotionCor2 with a 5 × 5 patch and a two-fold binning [89]. The CTF was estimated using CTFFIND4 [90] and corrected using CTFPHASEFLIP in IMOD [91]. Tilt series were aligned using gold fiducial alignment and tomograms were reconstructed by weighted back-projection in IMOD [91]. The six best tomograms were binned to 11.12 Å pixel size, and denoised using IsoNet [92]. From these six tomograms, 100 vesicle-bound ATG2A-WIPI4 complexes were identified. Their angles with respect to the vesicle membrane were estimated from manual identification of the coordinates for three points: the vesicle center, the point where ATG2A touches the membrane, and the distal end of the rod-like ATG2A density. The denoised tomograms were segmented using Amira (Thermo Fisher Scientific) for the representation of membranes and ATG2A-WIPI4 complexes.

### Pulldown assay

The purified ATG9A with C-terminal Twin-StrepII-6xHis tag and ATG2A with N-terminal Twin-StrepII-Flag tag were used for pulldown. 10 μM of full-length ATG2A and mini ATG2A (AA 1-443) was incubated with 1 μM of ATG9A in 200 uL reaction supplemented with 25 μL Nickel beads respectively. Control group containing 10 μM of full-length ATG2A and mini ATG2A (AA 1-443) was incubated with 25 μL Nickel beads respectively. After 2 hours incubation at 4 °C, beads were washed three times with 20 mM HEPES, pH 8.0, 150 mM NaCl, 1 mM TCEP, 0.09 mM LMNG supplemented with 20mM imidazole, and eluted with supplemented with 300 mM imidazole. The sample were detected by SDS–PAGE and western blotting with anti-Flag antibody. The pulldown was repeated three times with similar results. The quantification of SDS-PAGE band densitometry was measured by ImageJ. Data were analyzed by GraphPad Prism 6, and error bars represent SEM.

### Western blotting

The pulldown elution sample were further resolved on SDS-PAGE and followed by transfer onto a PVDF membrane (BIO-RAD). Then membranes were blocked in 5% milk/TBST for 2 hours at room temperature and incubated with primary antibodies at 4°C overnight. Following incubated with secondary antibodies conjugated to horseradish peroxidase (anti-mouse, Cwbio) for 2 hours at room temperature and visualized with BeyoECL Plus (Beyotime). The images were acquired using Amersham Imager 680 (GE Healthcare).

### Quantification and statistical analyses

Data were analyzed using GraphPad Prism 6 software. The statistical details are reported in material and methods and figure legends, including statistical analysis performed, error bars, and repeat numbers.

## Acknowledgements

The authors thank the cryo-EM (KEMC) and Advanced Mass Spectrometry Facility (KMS) of Kobilka Institute of Innovative Drug Discovery, the Chinese University of Hong Kong (Shenzhen) for the support. The cryo-ET data were collected at the Umeå Centre for Electron Microscopy (UCEM), a SciLifeLab National Cryo-EM facility and part of National Microscopy Infrastructure, NMI VR-RFI 2016-00968), supported by instrumentation grants from the Knut and Alice Wallenberg Foundation and the Kempe Foundations. We are grateful to Dr. Ming-Yuan Su for critically reading the manuscript. This work was supported by the National Natural Science Foundation of China NO. 31950410540 (G.S.) and 31971179 (L.Z.), Shenzhen Fundamental Research Fund JCYJ20200109150003938 (L.Z.) and RCYX20200714114645019 (L.Z.), the Foreign Young Talent Program NO. QN2021032004L (G.S.) and the Swedish Research Council (grants 2018–05851, 2021–01145, to L.A.C.). Y.W. and R.T. were supported by a Ganghong Young Scholar Development Fund at the Chinese University of Hong Kong, Shenzhen. The computational work was supported by the Warshel Institute for Computational Biology (funding from Shenzhen City and Longgang District), the fund from Shenzhen-Hong Kong Cooperation Zone for Technology and Innovation (HZQB-KCZYB-2020056).

## Contributions

Y. W., L.-A.C., L.Z. and G.S. designed the experiments. Y. W. cloned constructs, expressed and purified proteins, collected and processed electron microscopy data, built models, prepared liposome and performed lipids transfer activity assay. S.D. performed cryo-ET sample, collected and processed tomography data. R.T. analyzed molecular dynamics simulations. G.S. and X.M. performed XL-MS experiment. G.S., L.-A.C. and Y.W. wrote the manuscript. This work is supervised by L.Z., L.-A.C. and G.S.

---

**Fig. S1:**
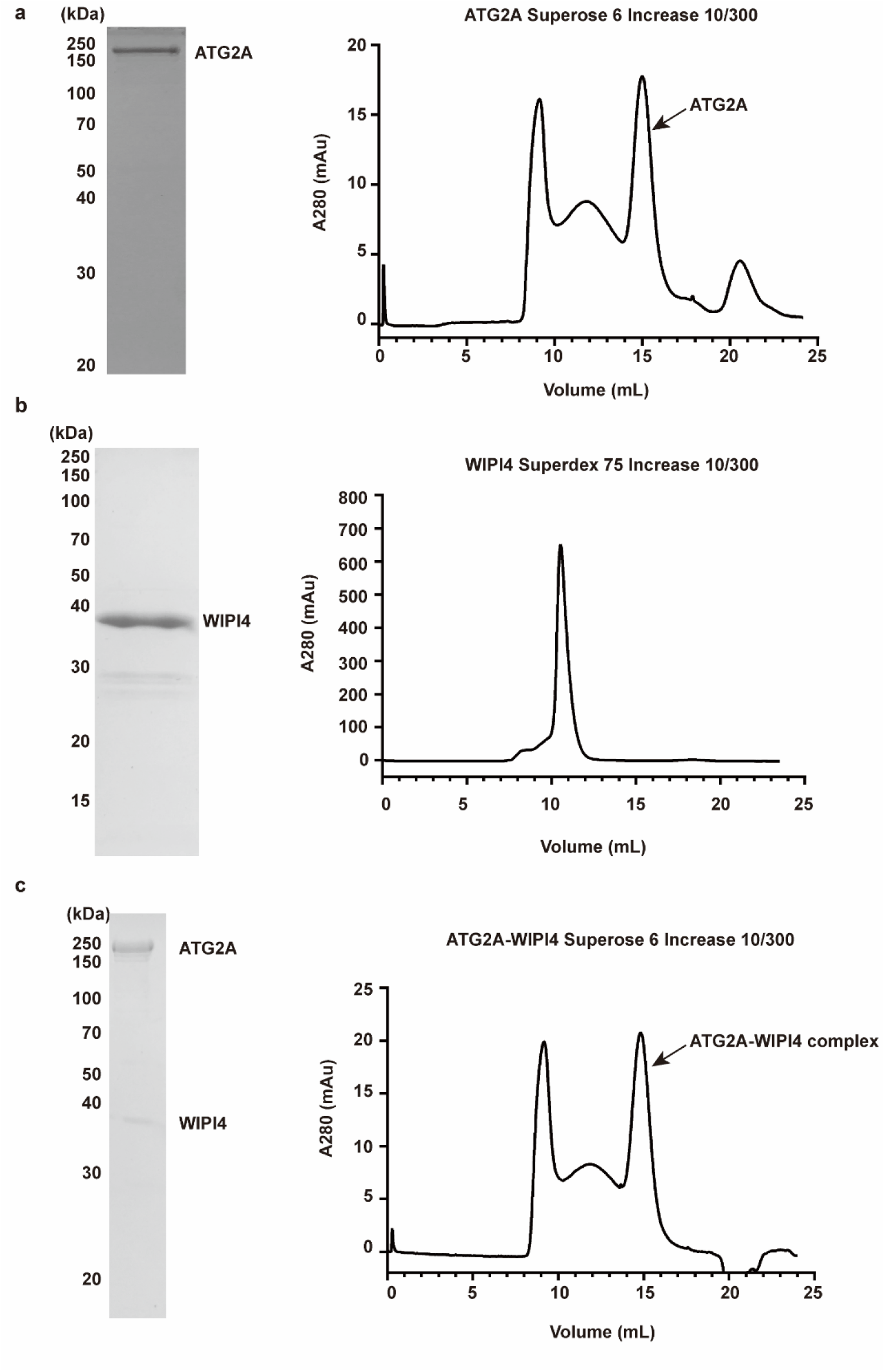
Protein purification SDS-PAGE results and gel filtration profiles. The SDS-PAGE (left) and gel filtration profile (right) of ATG2A alone (**a**), WIPI4 alone (**b**) and ATG2A-WIPI4 complex (**c**).

**Fig. S2:**
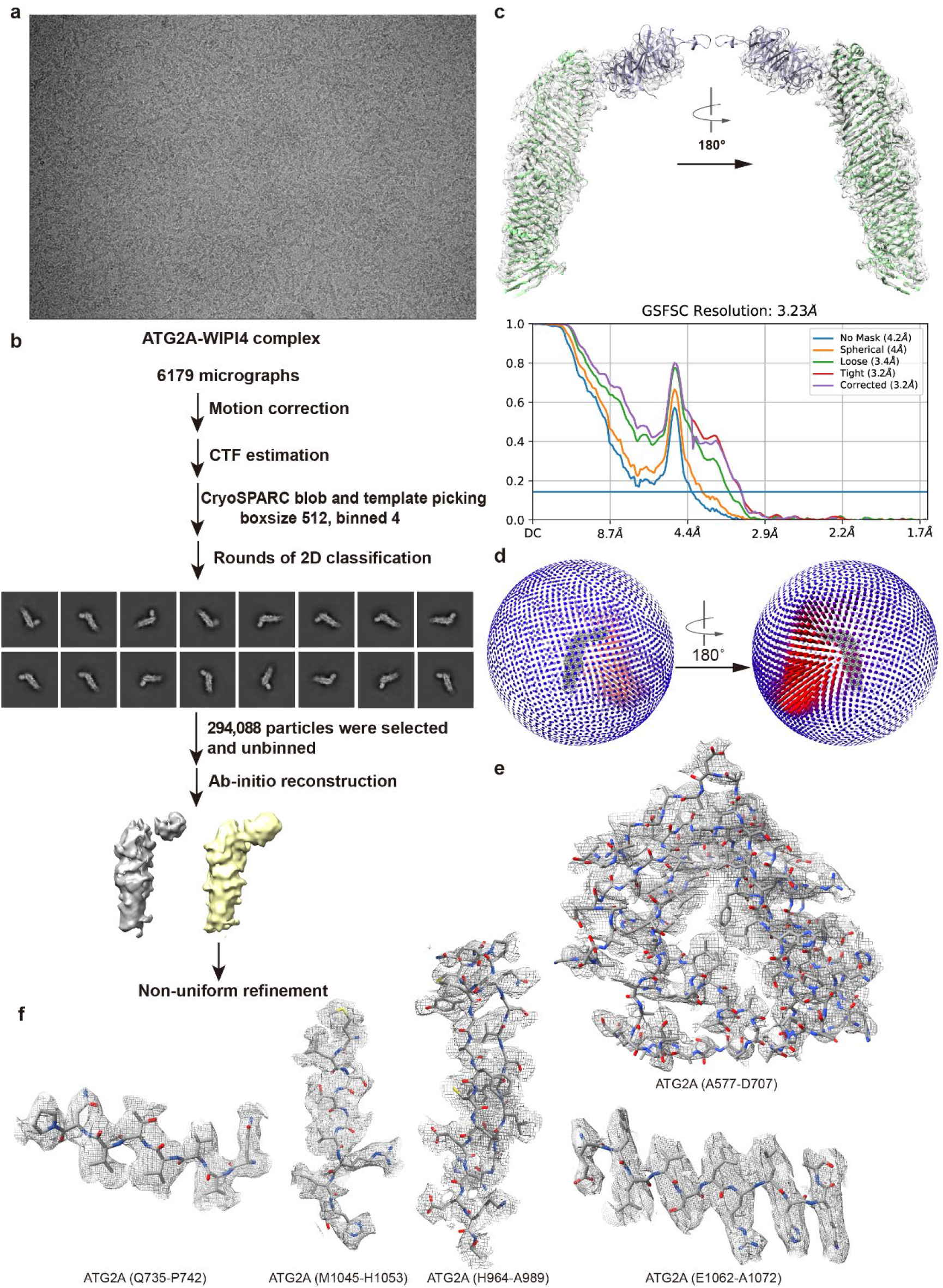
Imaging processing procedure of the ATG2A-WIPI4 complex. **a,** The representative micrograph after motion correction of ATG2A-WIPI4 complex. **b,** Cryo-EM data processing flow chart of ATG2A-WIPI4 complex. **c,** The ATG2A-WIPI4 model fitted into the density map. **d,** Particle orientation distribution of ATG2A-WIPI4 map. **e,** The density map with corresponding structural model of ATG2A A577-D707, relative to the region shown in Fig 2b. **f**, The representative density map with corresponding structural model of ATG2A.

**Fig. S3:**
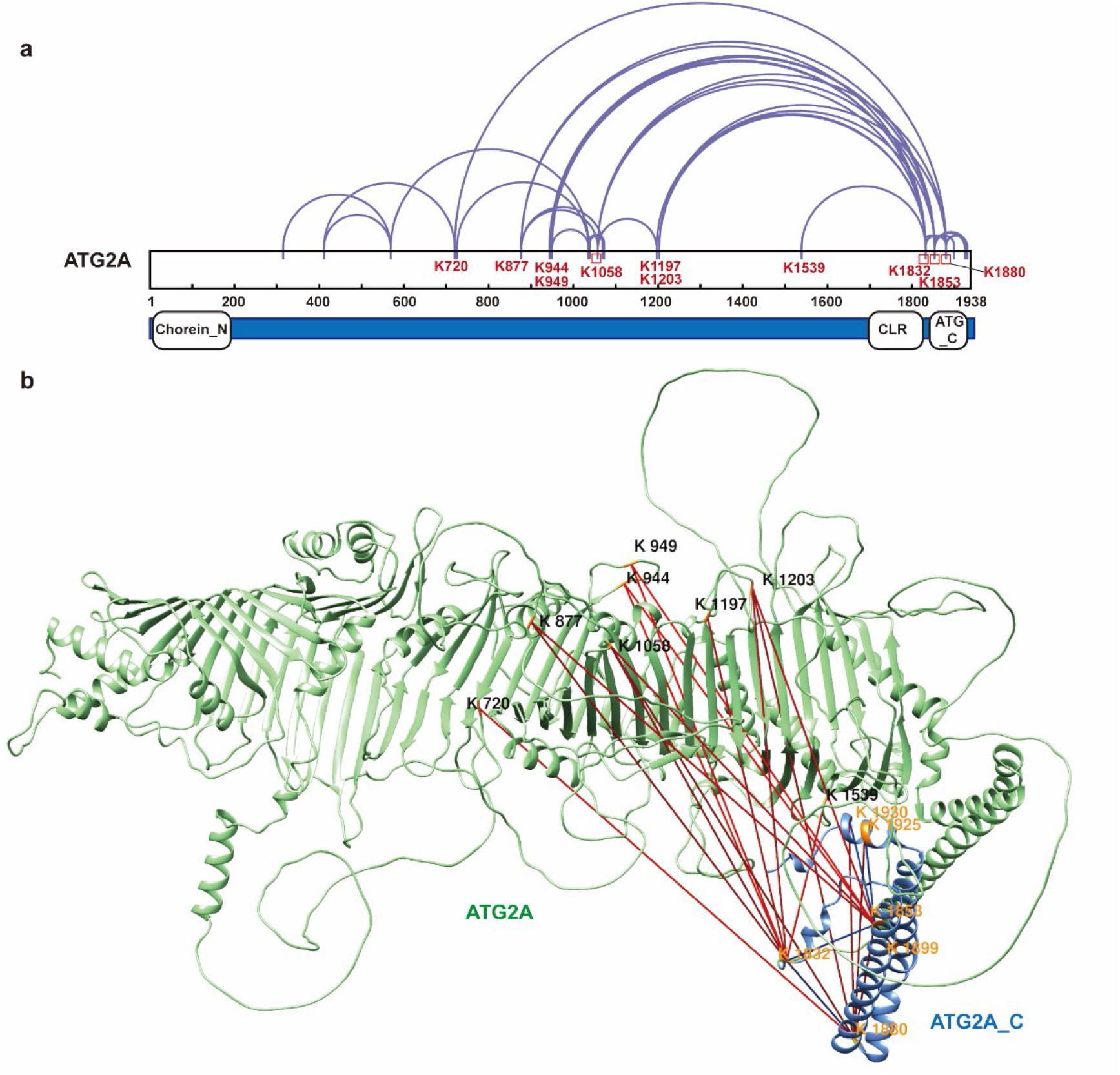
ATG2A intramolecular crosslinks. **a,** Linear plot displaying all the identified crosslink sites for ATG2A. **b,** Intramolecular ATG2A crosslinks mapped onto AlphFold model. ATG2A_C region (blue) crosslinks with ATG2A rod-like domain (green). Unsatisfied (>33 Å) and satisfied crosslinks are indicated in red and blue, respectively.

**Fig. S4:**
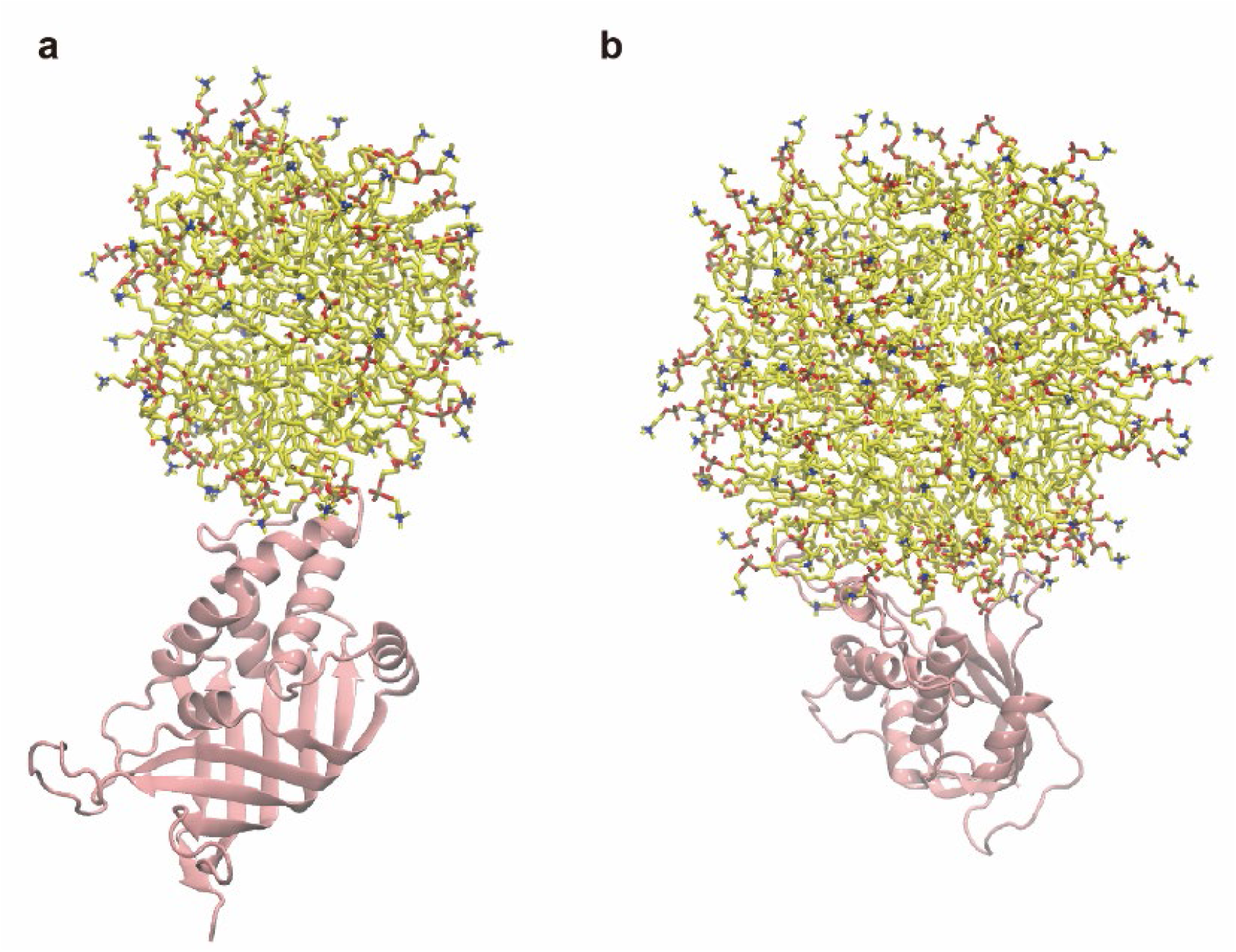
The different binding poses between *sp*Atg2_NR_ and DOPC micelle in simulations. **a,** The DOPC micelle contacts the lateral helix of *sp*Atg2^NR^. *sp*Atg2^NR^ is colored in pink and the DOPC molecules are colored in yellow. **b,** The DOPC micelle contacts the edge of the folded sheet of *sp*Atg2^NR^.

**Fig. S5:**
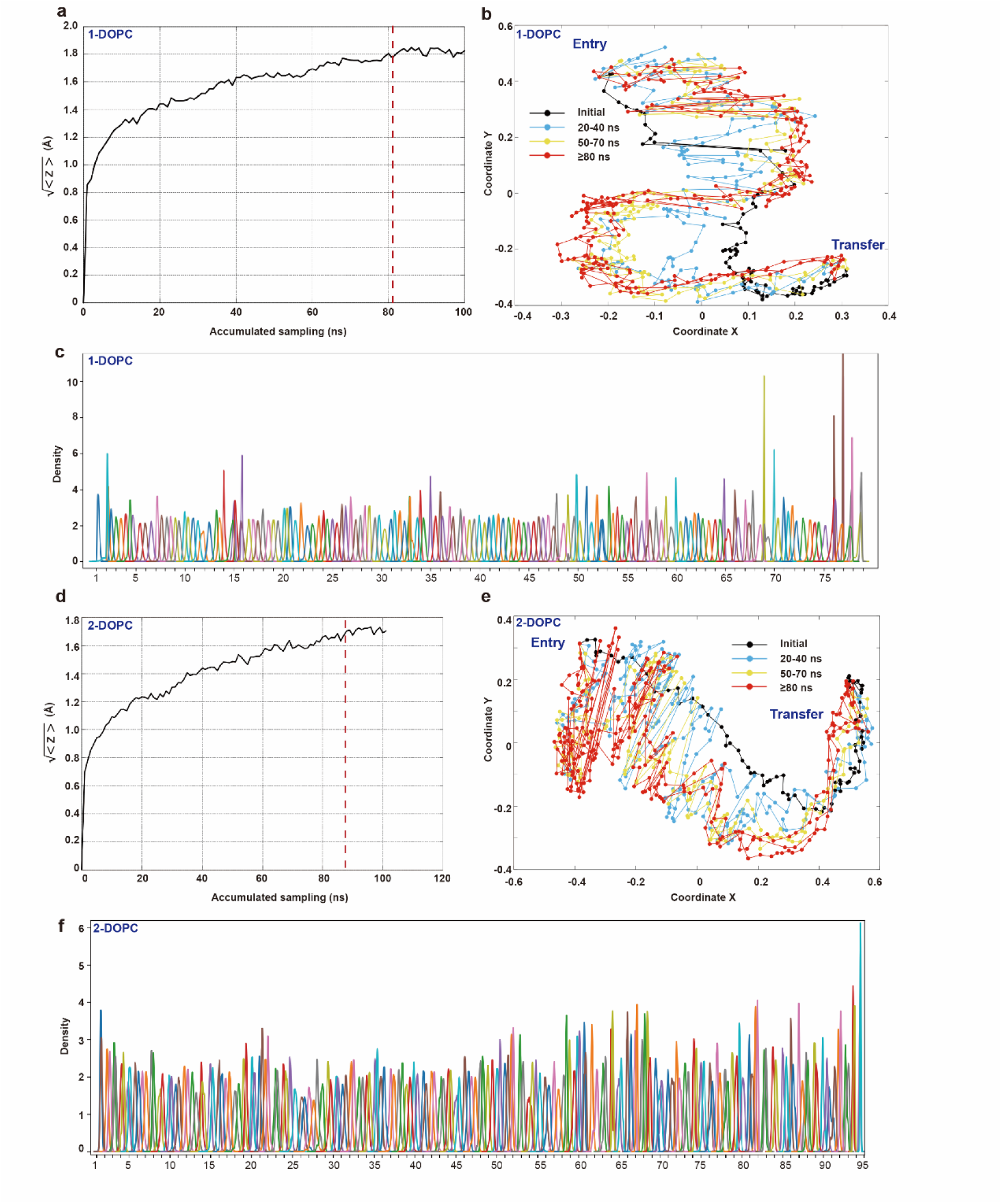
Convergence check and sample distribution of 1-DOPC and 2-DOPC simulations. **a, d,** Convergence of the optimization processes for the path of 1-DOPC **(a)** and 2-DOPC **(d)** transfer by ATG2A^NR^ is measured by the progress of ⟨z⟩ along the accumulated sampling time. The initial path is the reference. Convergence time is highlighted by a red dashed line. **b**, **e** The paths at different TAPS iterations are mapped to a two-dimensional space by multidimensional scaling (MDS) for 1-DOPC **(b)** and 2-DOPC **(e)**. The accumulated sampling time for each iteration is highlighted by different colors. **c, f,** Sample distribution in all windows of the umbrella sampling along the PCV_s for the path of lipid transfer by ATG2A^NR^ for 1-DOPC **(c)** and 2-DOPC **(f)**.

**Fig. S6:**
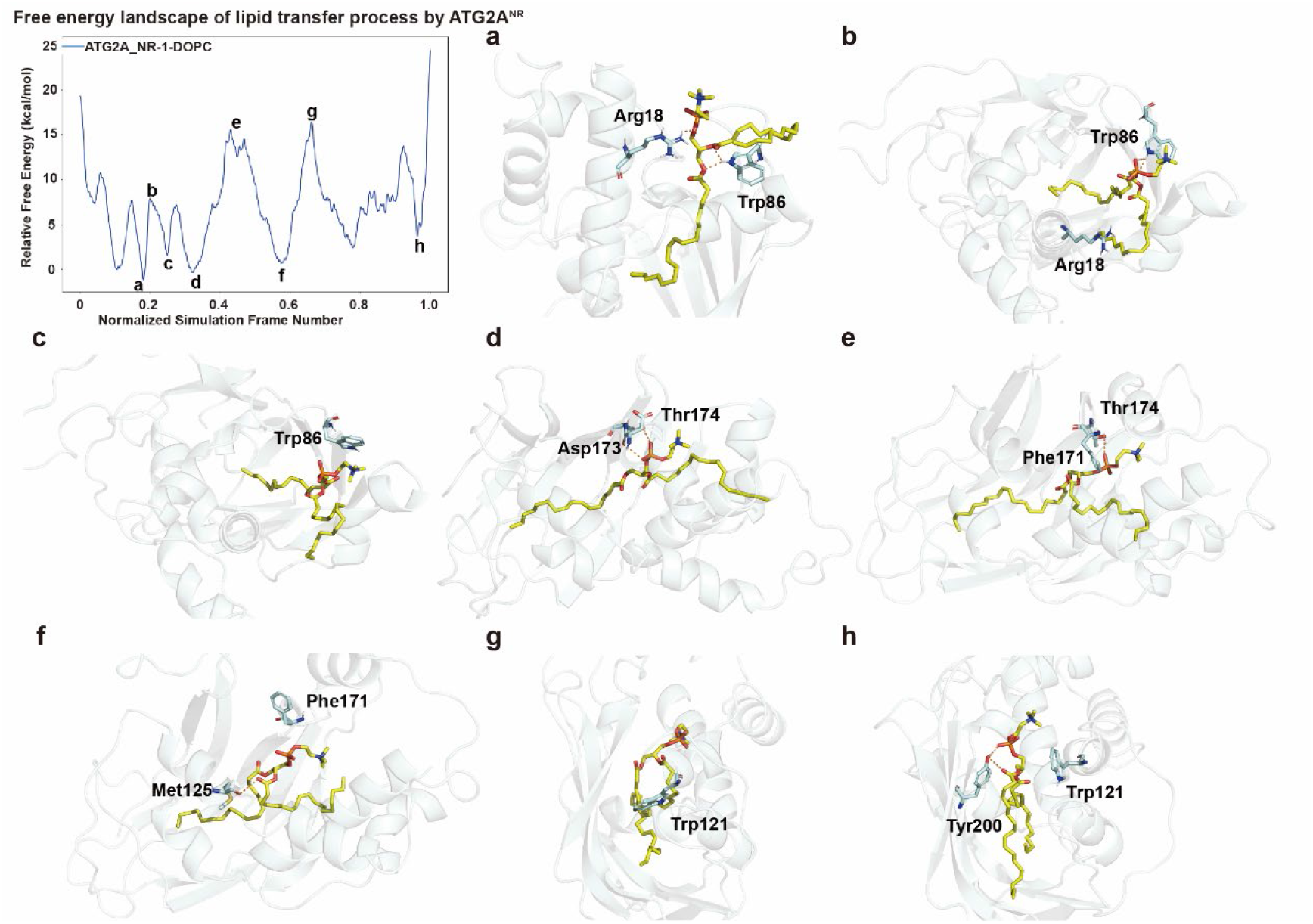
The detailed process of lipid transport by ATG2A_NR_. The free energy curve of 1-DOPC simulation is drawn in blue. **a-h** are the structures corresponding to the points marked on the curve. ATG2A^NR^ is colored in pale cyan and the DOPC is colored in yellow.

**Fig. S7:**
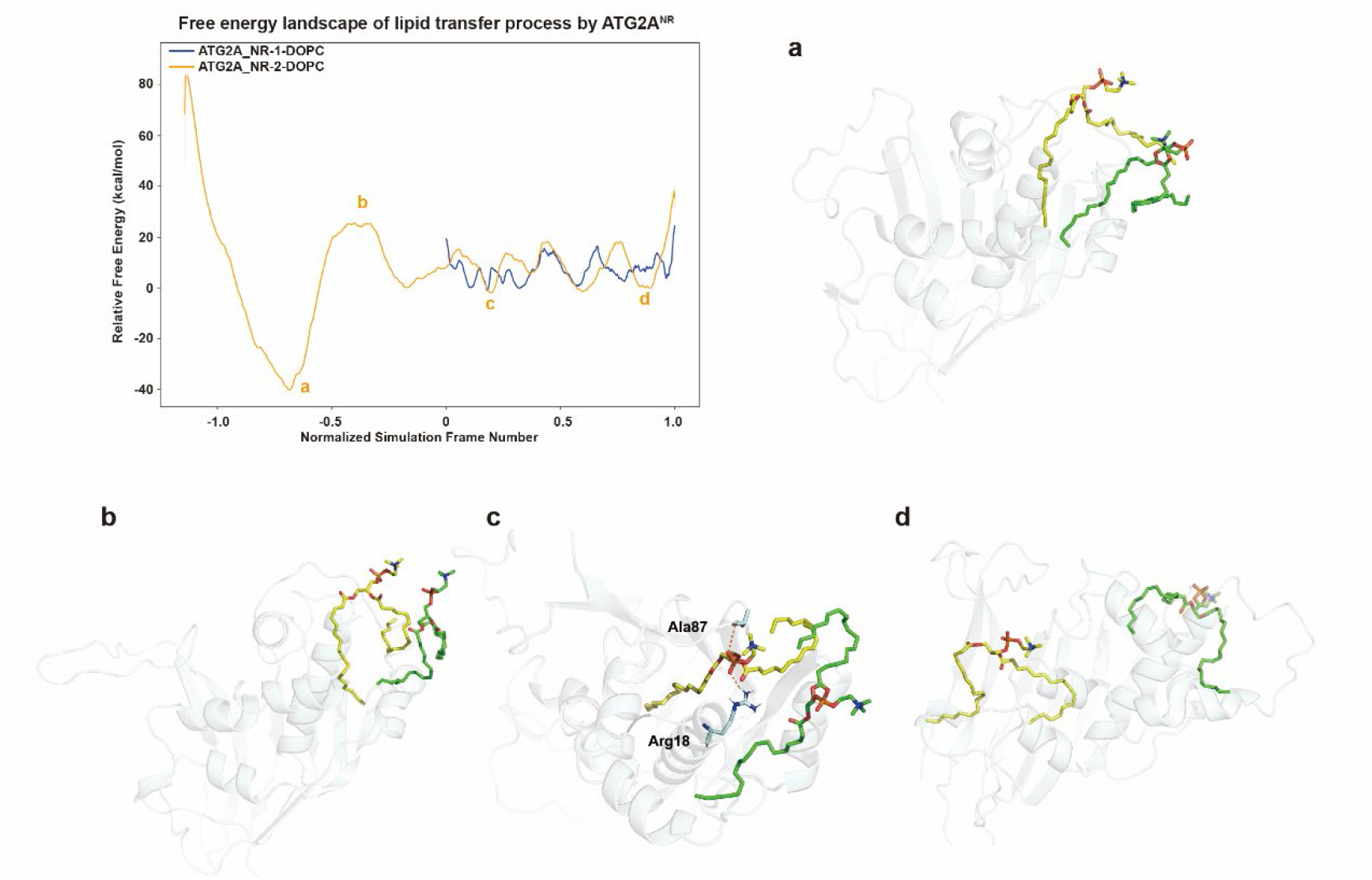
The comparison of 1-DOPC and 2-DOPC simulations of lipid transporting process by ATG2A_NR_. The free energy curves of 1-DOPC and 2-DOPC simulations are drawn together. The one of 1-DOPC system is colored in blue and the one of 2-DOPC system is colored in orange. The two curves are aligned according to conformational changes. **a-d** are the structures corresponding to the points marked on the curve. ATG2A^NR^ is colored in pale cyan. The entering DOPC is colored in yellow and the one staying outside in green.

**Fig. S8:**
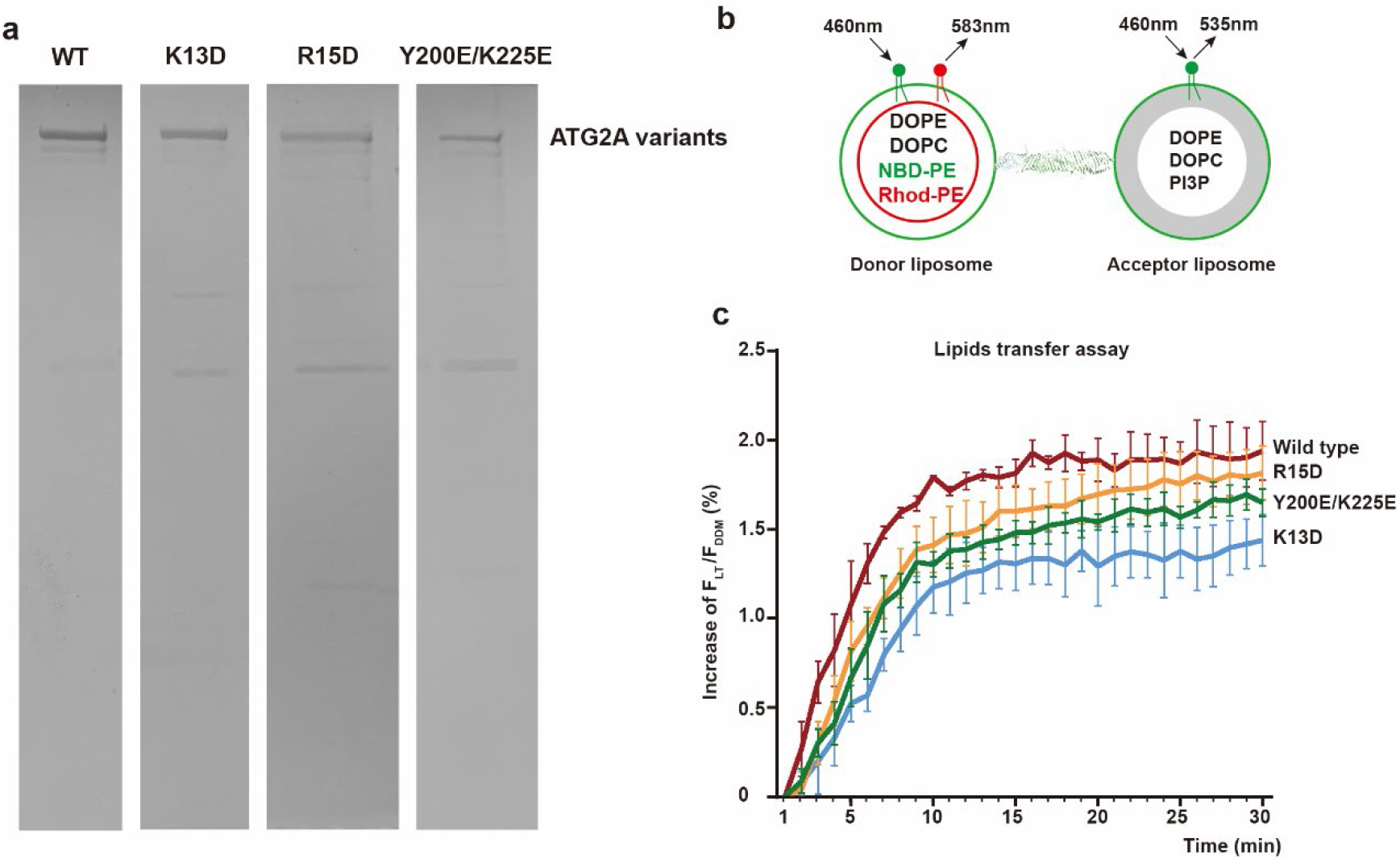
Two-step purified ATG2A wild type and mutants and activity assay. **a,** ATG2A wildtype (WT) and mutants K13D, R15D, Y200E/K225E were purified by GST and Strep-Tactin affinity chromatography. The purified proteins were used for lipid transfer activity assay. **b,** The activity assay model of ATG2A transfer lipids *in vitro*. **c,**The comparison of lipids transfer activity between wild type and mutants. The lipid transfer activity is measured by the NBD fluorescence which is not quenched by Rhodamine. F_LT_ means the NBD fluorescence intensity at each time point, followed by treatment with 0.4% DDM for 40 min and denoted as F_DDM_, these two values are measured at 535 nm. The increase in F_LT_/F_DDM_ was calculated by subtracting F_LT_/F_DDM_ of control group from protein-containing group, and minus the value at 1 min point. And the curve shows the mean values from three independent experiments.

**Fig. S9:**
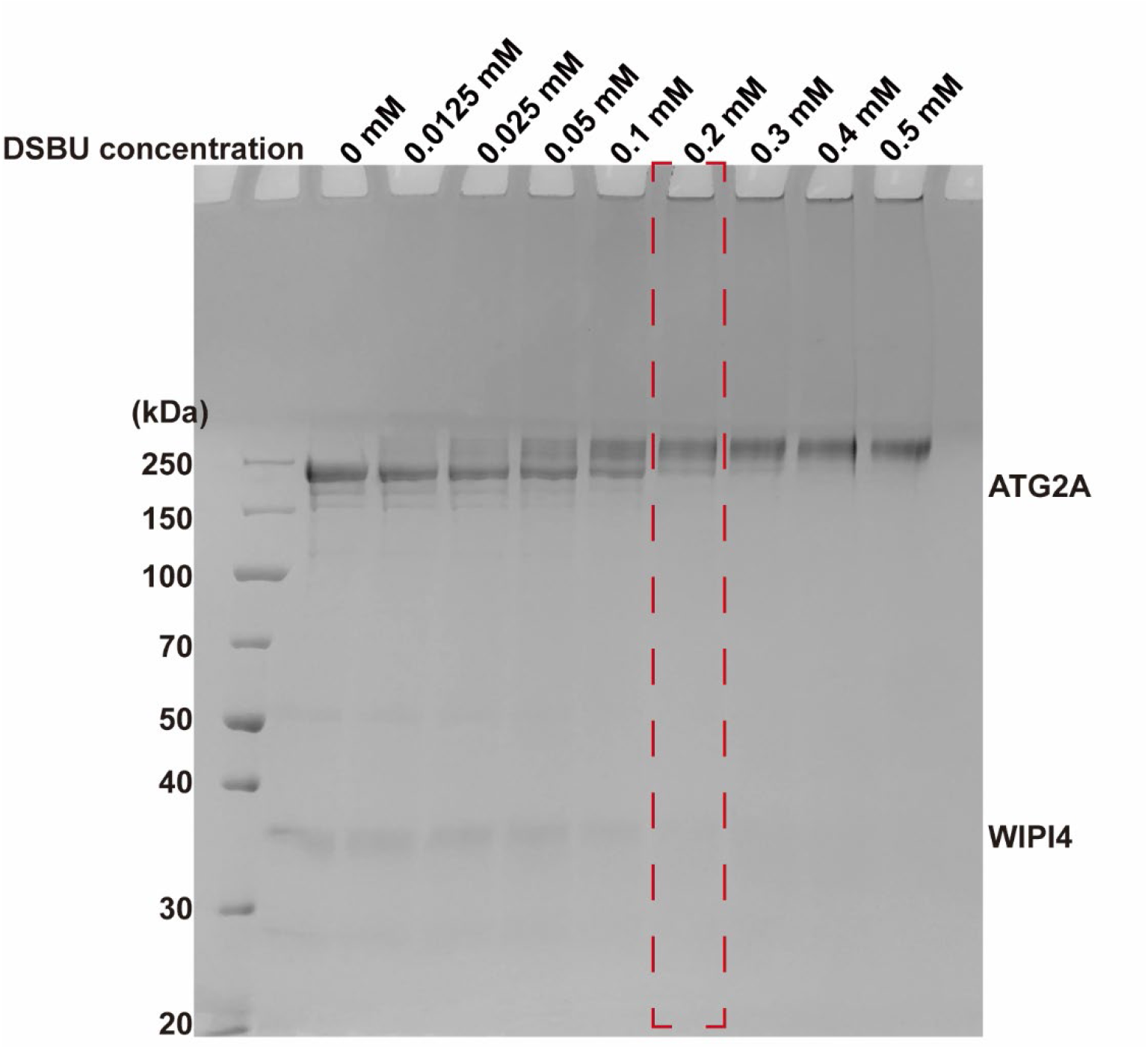
DSBU crosslinking. DSBU was used to crosslink *in vitro* reconstituted ATG2A-WIPI4 complex. A series of DSBU concentration were tested at RT, and 0.2 mM crosslinking condition (red dash line) was selected for mass spectrometry analysis.

**Fig. S10:**
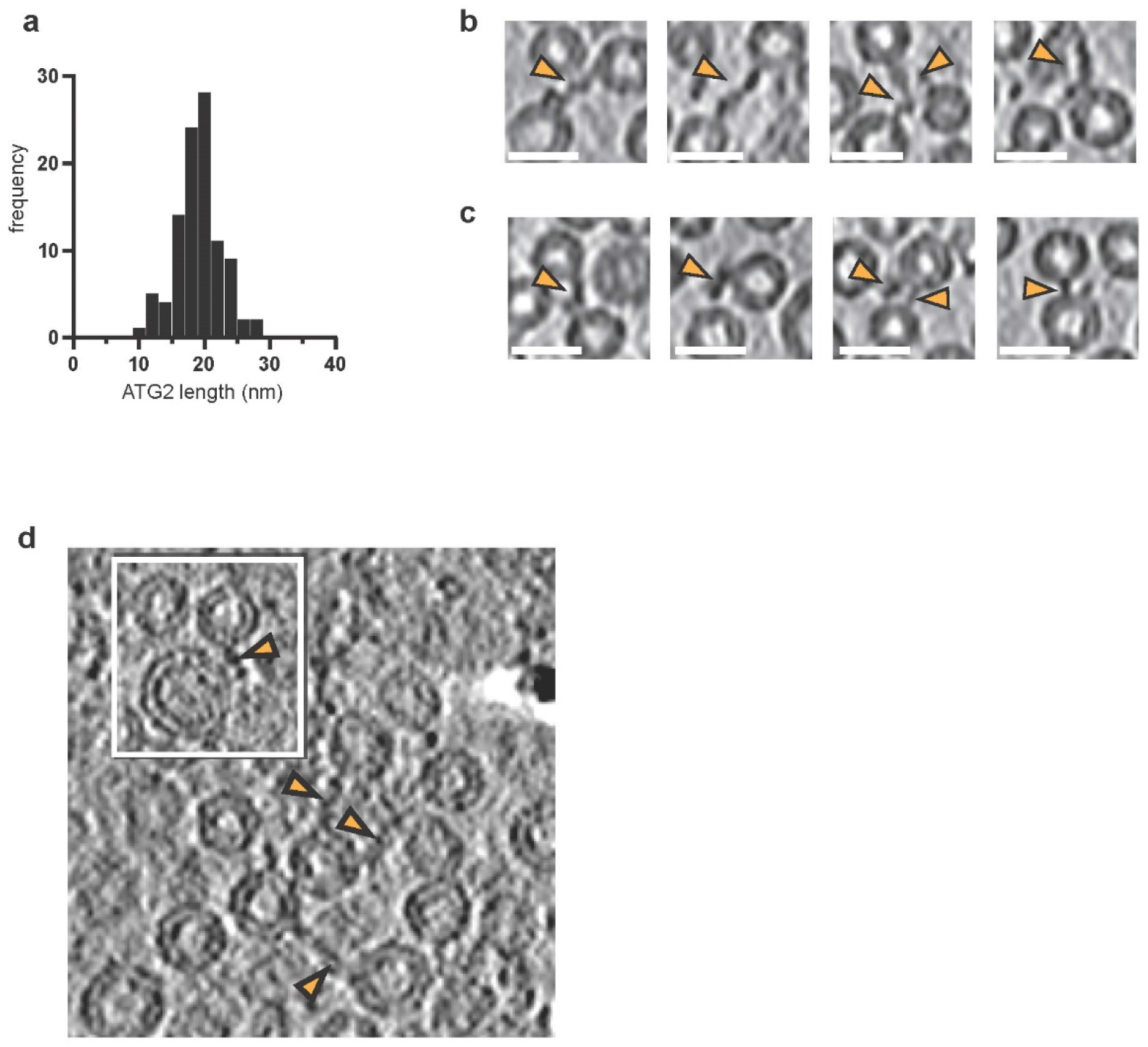
Analysis of the ATG2A-WIPI4 membrane tethering by cryo-electron tomography. **a,** Distribution of lengths of the ATG2A-WIPI4 complexes tethering SUVs in cryo-tomograms. The average length is 19.0 nm (SD=3.5, N=100). (**b-c**) Crops of cryo-electron tomograms showing the different orientations that ATG2A-WIPI4 adopts on the membranes. ATG2A-WIPI4 associates orthogonally (**b**), and at different angles to the tethered membranes (**c**). Scale bar 20 nm. **(d)** Same computational slice as shown in Fig. 5a, but through an unfiltered version of the same tomogram. Arrowheads indicate same protein densities as in Fig. 5a. The black density in the upper right periphery is a fiducial gold particle.

**Fig. S11:**
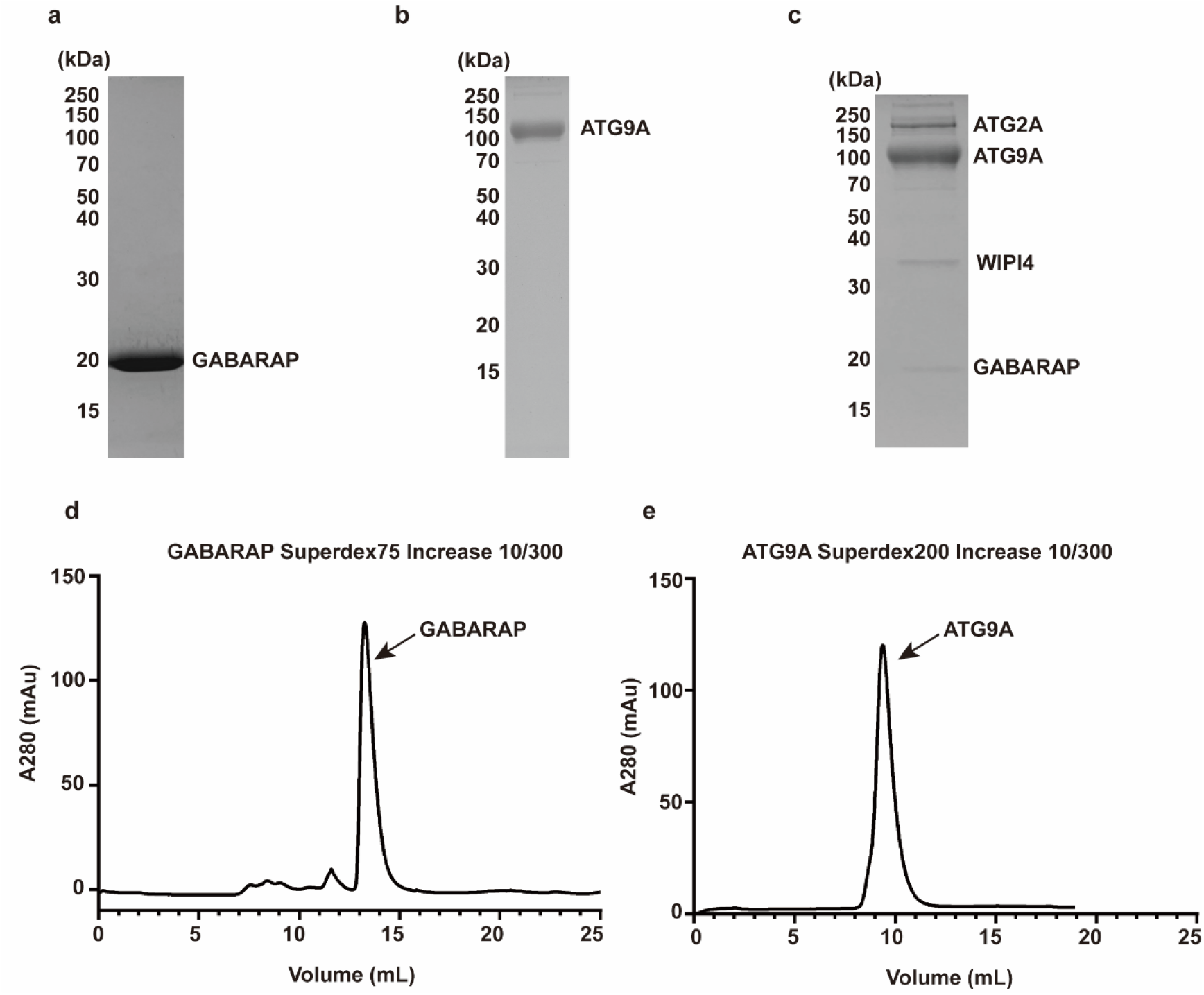
Purification and in vitro reconstitution of the ATG2A-WIPI4-GABARAP-ATG9A complex. The SDS-PAGE of GABARAP (**a**), ATG9A (**b**), and reconstituted ATG2A-WIPI4-GABARAP-ATG9A complex (**c**). Gel filtration profile of GABARAP (**d**) and ATG9A (**e**).

**Fig. S12:**
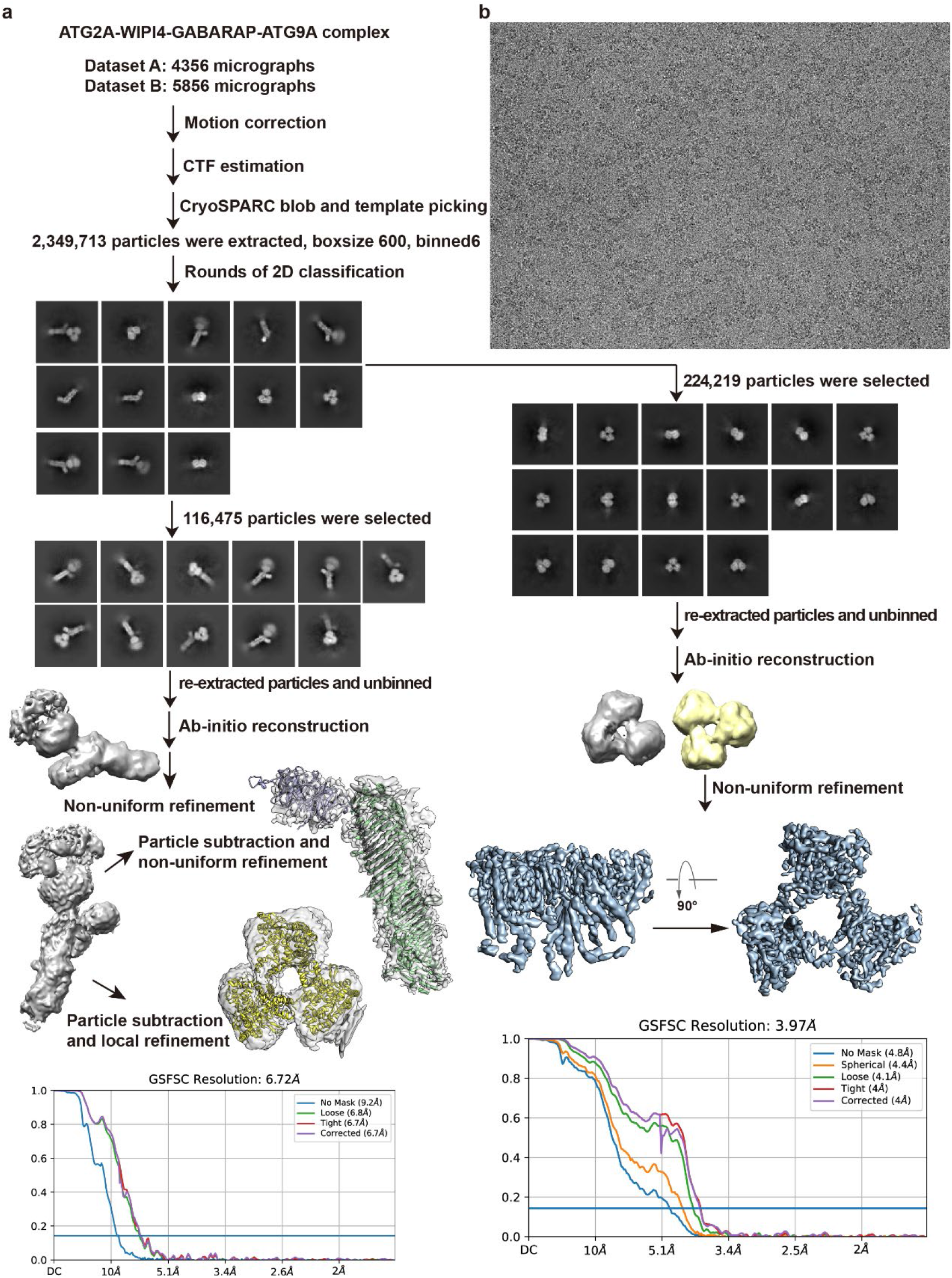
Imaging processing procedure of the C-terminally bound ATG2A/WIPI4-ATG9A complex. **a,** Imaging processing workflow of ATG2A/WIPI4-ATG9A complex and dissociated ATG9A for refinement and reconstruction of density maps. **b,** A representative cryo EM micrograph of ATG2A/WIPI4-ATG9A complex.

**Fig.S13:**
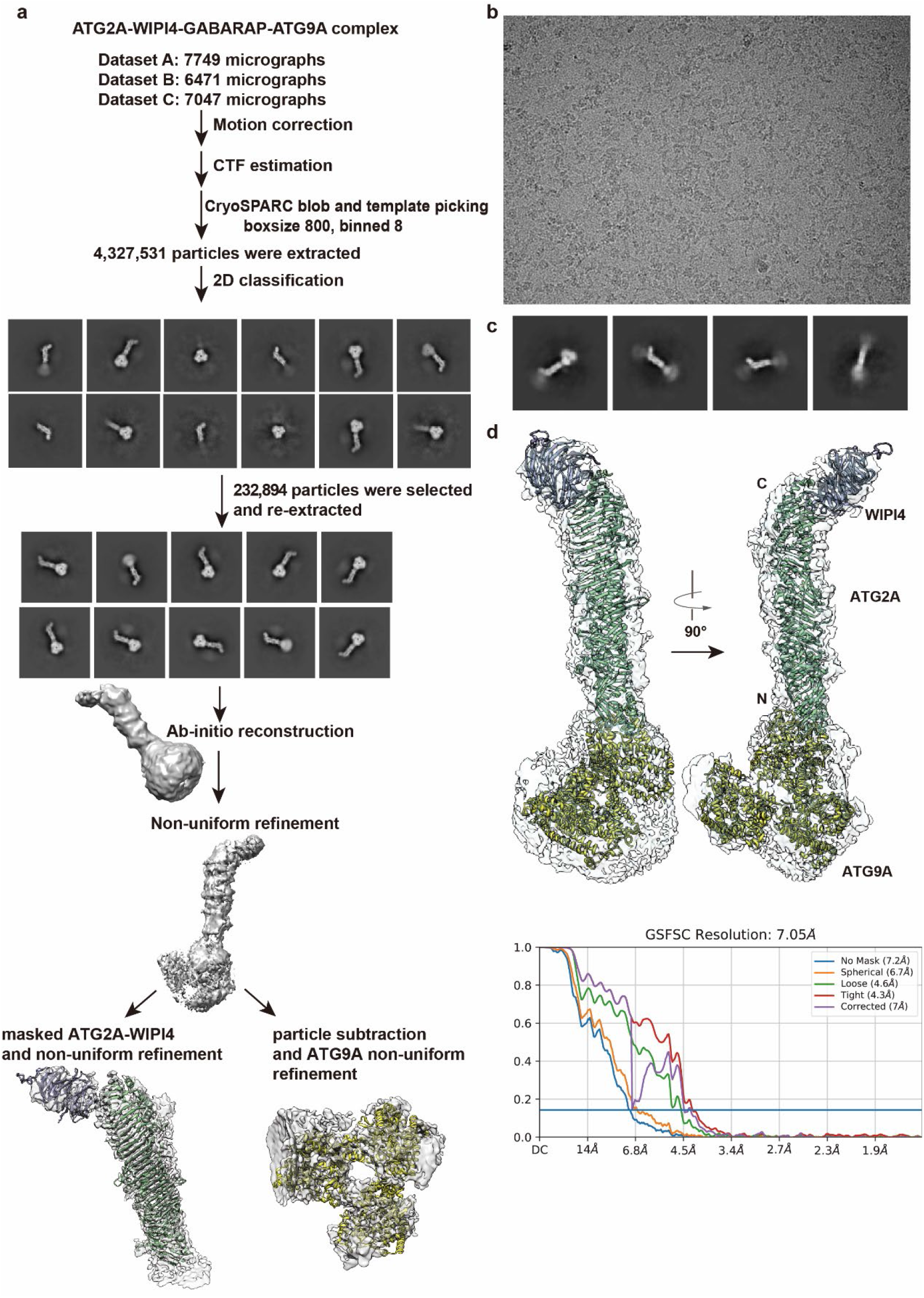
Imaging processing procedure of the N-terminally bound ATG2A/WIPI4-ATG9A complex. **a**, Imaging processing workflow of N-terminally bound ATG2A/WIPI4-ATG9A complex. **b**, A representative cryo-EM micrograph of ATG2A/WIPI4-ATG9A complex. **c**, Representative 2D class averages indicating that ATG9A flexibly binds to both N- and C-terminus of ATG2A. **d**, Model fitting of ATG9A and ATG2A-WIPI4 into the low-resolution density map.

**Fig.S14:**
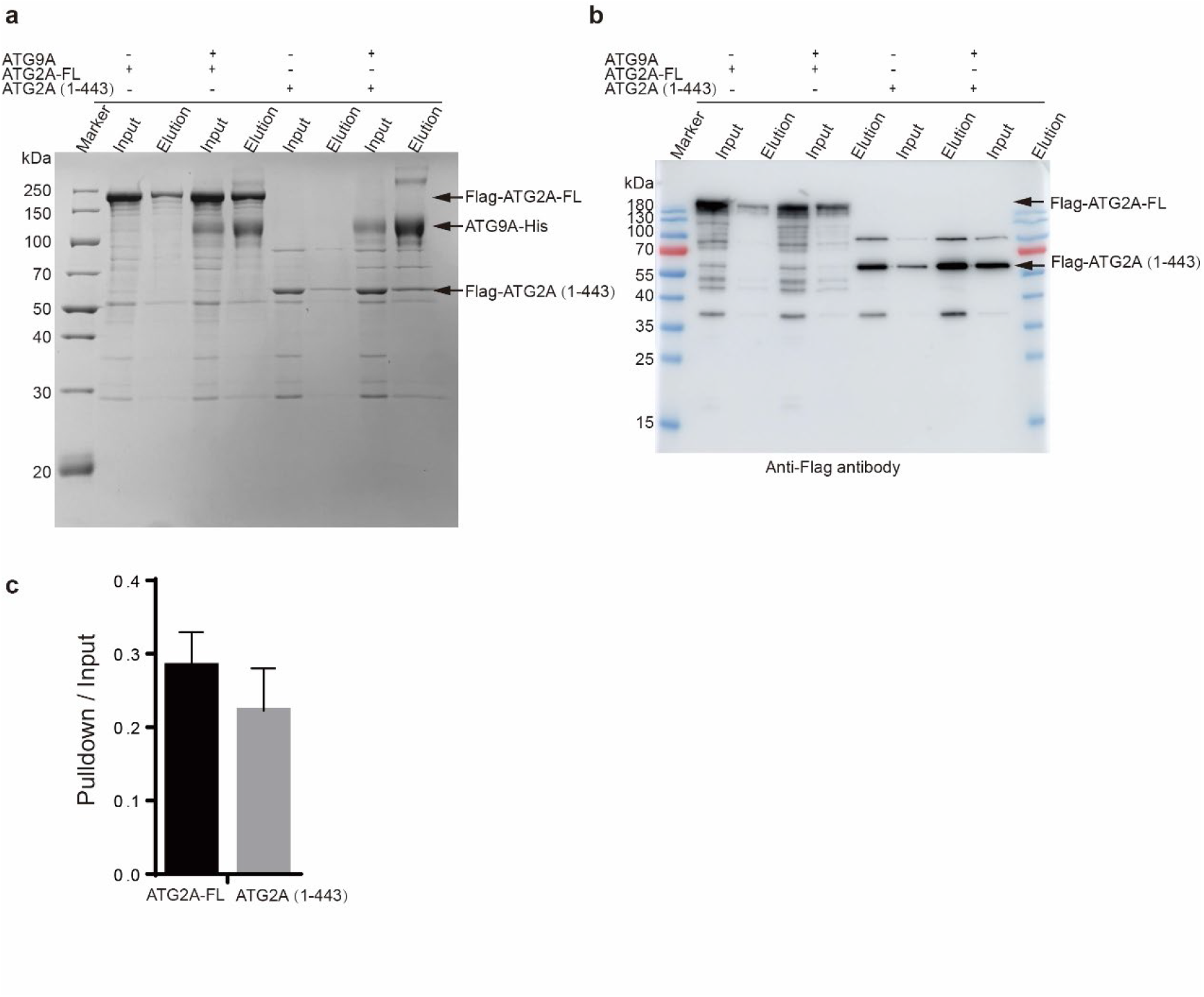
The interaction between N-terminal mini ATG2A and ATG9A complex. **a**, *In vitro* pull-down experiment of the full length ATG2A and N-terminal mini ATG2A (AA 1-443) interact with ATG9A. The experiment was repeated three times and visualized by SDS-PAGE. **b**, Western blotting of the pulldown results detected by anti-flag antibody. **c,** The quantification analysis of SDS-PAGE **(a)** to show the interaction between full-length ATG2A and mini ATG2A (AA 1-443) with ATG9A. Vertical coordinates is calculated by the mean density of elution band divided by input and subtract the control group.

**Fig. S15:**
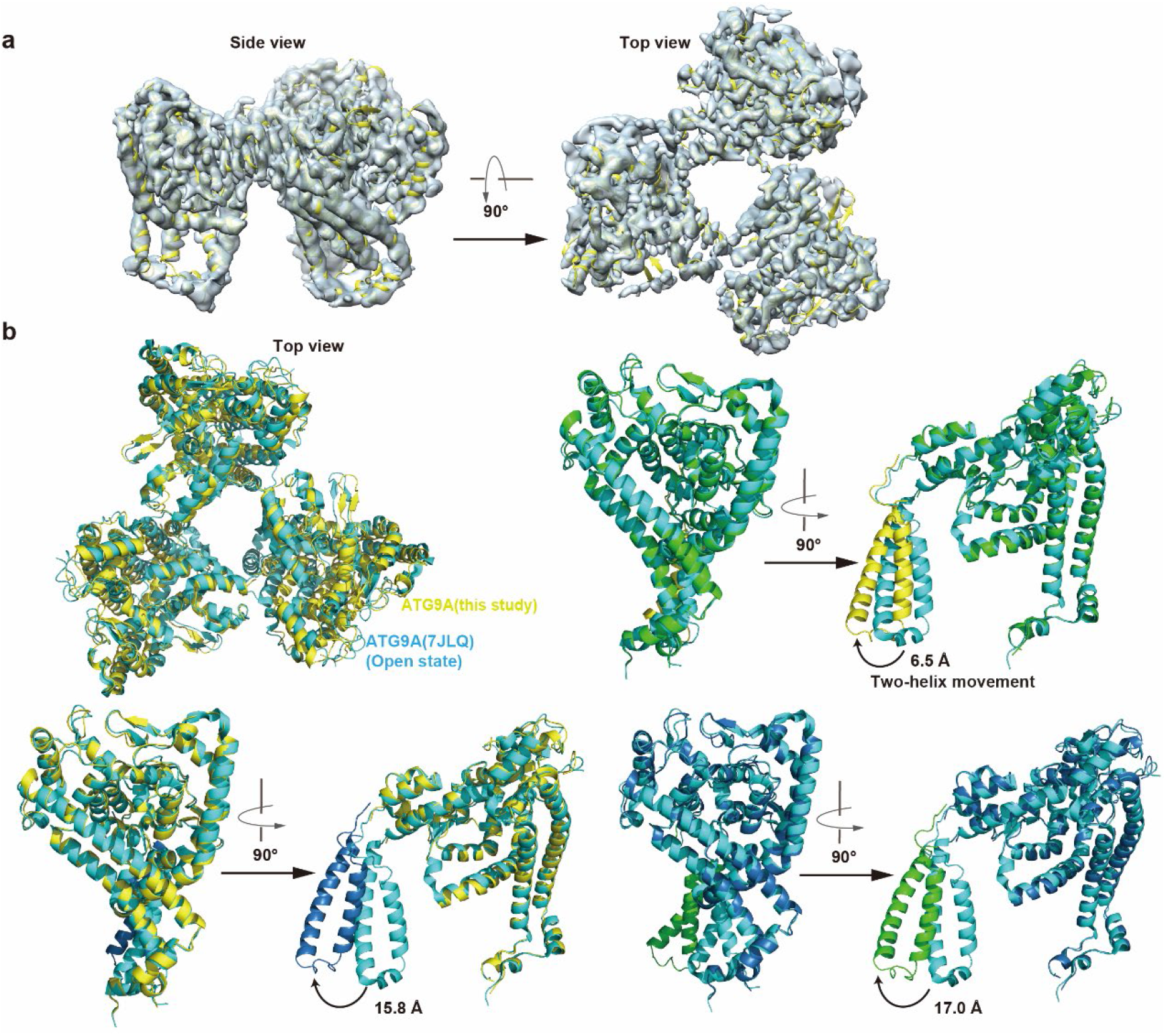
Structure of ATG9A in the open state. **a,** ATG9A model fitted into the density map. **b,** The comparison of ATG9A model in this study (yellow) with the published ATG9A model (7JLQ) (cyan) [24]. The superimposed cartoon of trimer (left) and protomer. The two-helix movement of ATG9A protomers was compared by measuring the distance between T409 in each protomer. The ATG9A promoter in this study is colored in yellow, green and sky-blue, and ATG9A model (7JLQ) was colored in cyan.

**Extended Table 1:**
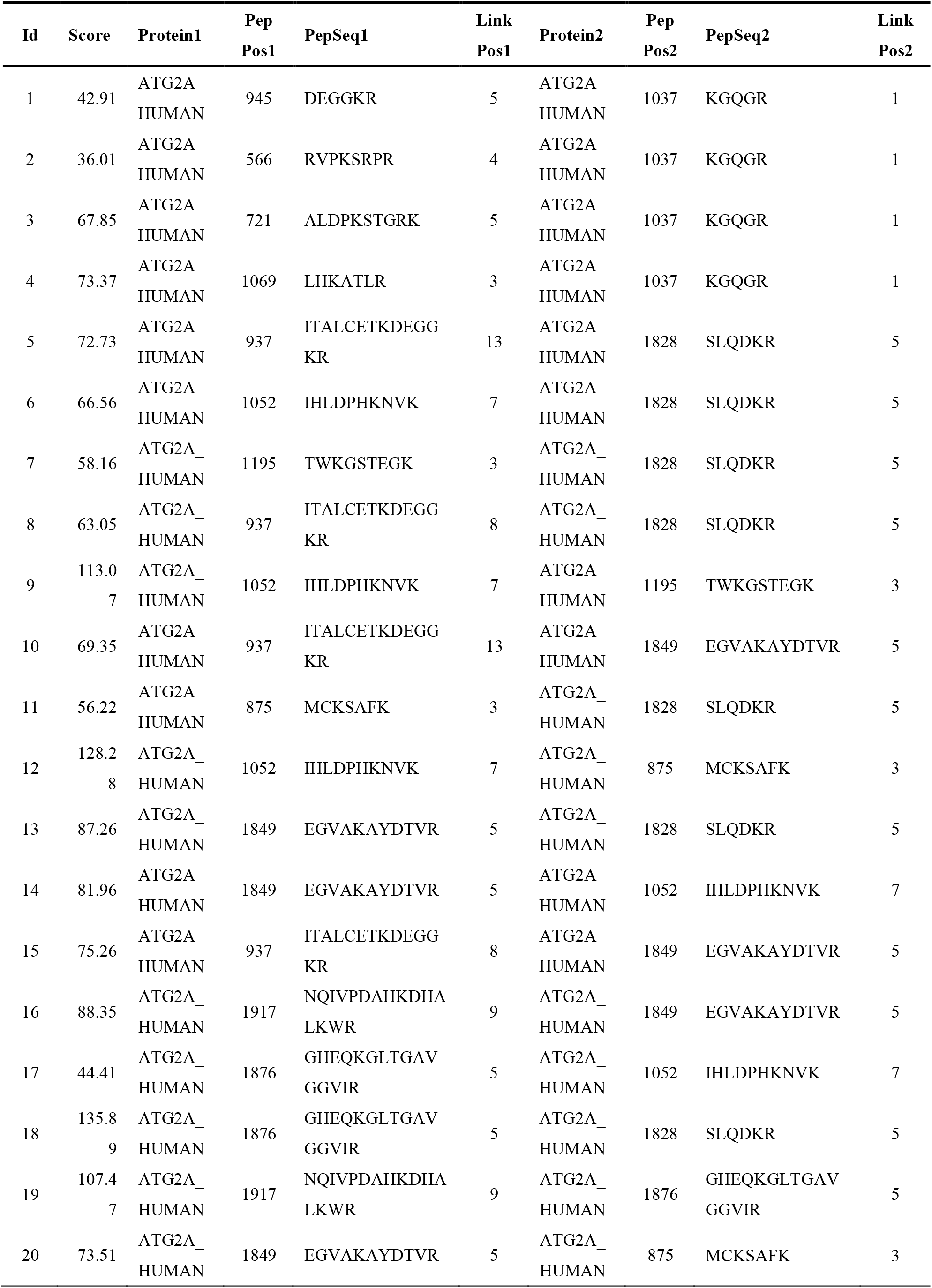
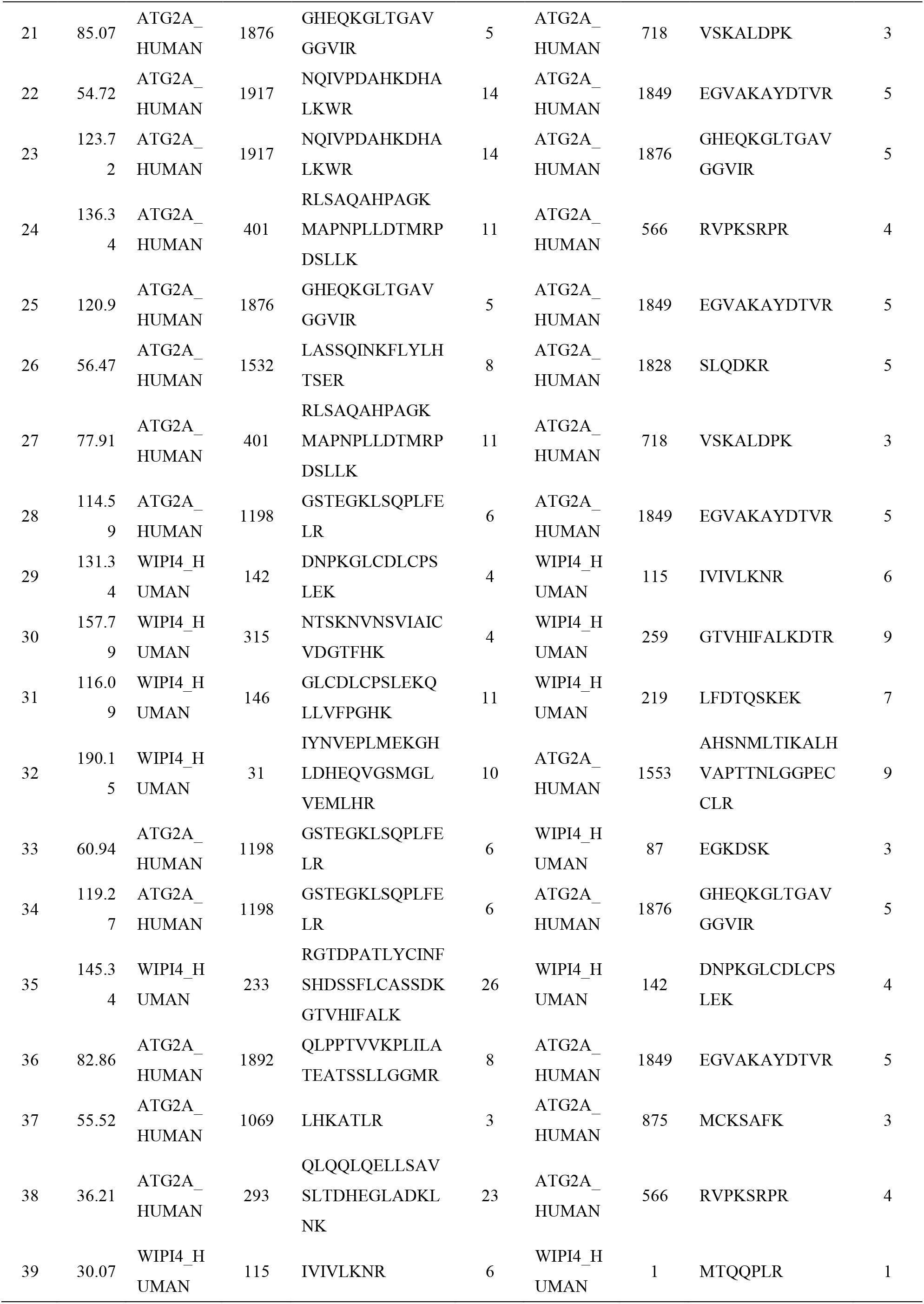
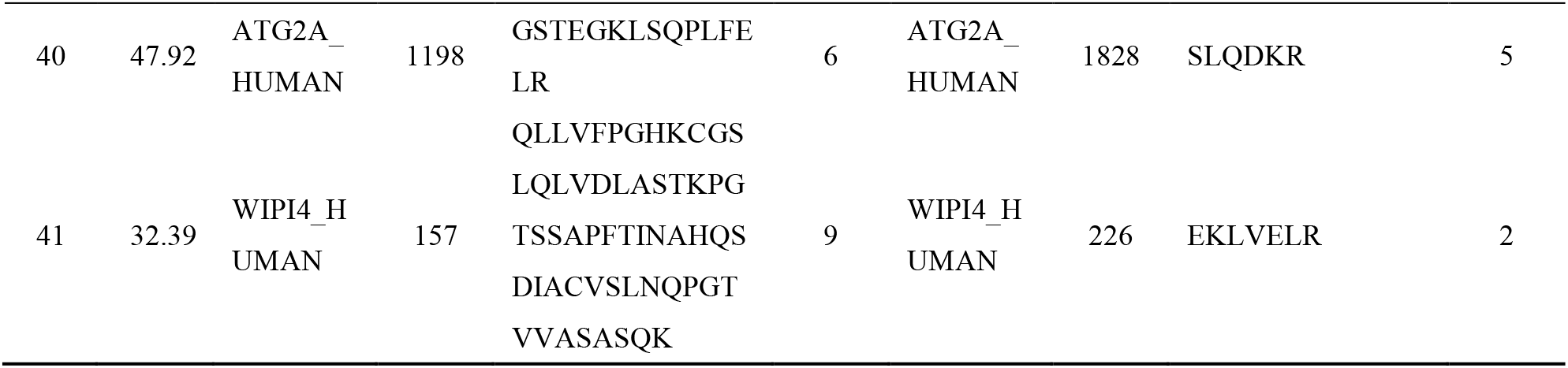
XL-MS results of ATG2A-WIPI4 complex.

**Extended Table 2:** The parameters in TAPS optimization.

|  |  |  |
| --- | --- | --- |
| System |  | ATG2A<sup>NR</sup>, 1DOPC/2DOPC |
| Sampling Time of Each Iteration |  | 1000 ps |
| Temperature |  | 310 K |
| Atom Sets | Alignment | C-alpha atoms of α-helices/β-sheets of ATG2A<sup>NR</sup> |
|  | RMSD | Heavy atoms of DOPC and residue 11-28, 35-39, 47-53, 56-61, 72-83, 94-99, 102-105, 120-125, 134-140, 159-166, 174-184, 193-201, 226-234 of ATG2A<sup>NR</sup> |
| Tolerant Distance for Neighbor Nodes |  | 1.3 Å |
| Well-tempered<br>Metadynamics<br>Simulation | Gaussian Height | 0.25 kJ/mol |
|  | Gaussian Width | 0.50 |
|  | Bias Factor | 10 |
|  | Length of tMD | 100 ps |
| Constant Force of tMD |  | 100000 kJ/mol/nm<sup>2</sup> |

**Video 1: The simulation of lipid extraction from DOPC micelle by *sp*Atg2_NR_.**

*sp*Atg2^NR^ is colored in pink, and DOPC in yellow. *sp*Atg2^NR^ is able to spontaneously absorb one phospholipid acyl chain to the entry of the lipid-transfer channel located on its N-terminal helical region.

**Video 2: The simulation of ATG2A_NR_ interaction with DOPC.**

ATG2A^NR^ is colored in cyan, and DOPC in yellow. ATG2A^NR^ can stably interact with DOPC acyl chain, similar to *sp*Atg2^NR^.

**Video 3: The simulation of lipid transfer process by ATG2A_NR_.**

ATG2A^NR^ is colored in cyan. The simulated lipid transfer process containing the entry, flip and transfer states, related to Fig. 3d-e.

**Video 4: The simulation of ATG2A_NR_ interaction with 2-DOPC.**

ATG2A^NR^ is colored in pale cyan, the entering DOPC is colored in yellow and the outside one in green. After the precursor molecule enters the protein, one acyl chains of the outside DOPC will occupies the entrance, related to Fig. S7.

